# Signal sequences target enzymes and structural proteins to bacterial microcompartments and are critical for microcompartment formation

**DOI:** 10.1101/2024.09.25.615066

**Authors:** Elizabeth R. Johnson, Nolan W. Kennedy, Carolyn. E. Mills, Shiqi Liang, Swetha Chandrasekar, Taylor M. Nichols, Grant A. Rybnicky, Danielle Tullman-Ercek

**Affiliations:** Department of Chemical and Biological Engineering, Northwestern University, Evanston, Illinois, USA; Interdisciplinary Biological Sciences Program, Northwestern University, Evanston, Illinois, USA; Department of Chemistry and Biochemistry, Loyola University Chicago, Chicago, Illinois, USA; Chemistry of Life Processes Institute, Northwestern University, Evanston, Illinois, USA; Center for Synthetic Biology, Northwestern University, Evanston, Illinois, USA

## Abstract

Spatial organization of pathway enzymes has emerged as a promising tool to address several challenges in metabolic engineering, such as flux imbalances and off-target product formation. Bacterial microcompartments (MCPs) are a spatial organization strategy used natively by many bacteria to encapsulate metabolic pathways that produce toxic, volatile intermediates. Several recent studies have focused on engineering MCPs to encapsulate heterologous pathways of interest, but how this engineering affects MCP assembly and function is poorly understood. In this study, we investigated the role of signal sequences, short domains that target proteins to the MCP core, in the assembly of 1,2-propanediol utilization (Pdu) MCPs. We characterized two novel Pdu signal sequences on the structural proteins PduM and PduB, which constitutes the first report of metabolosome signal sequences on structural proteins rather than enzymes. We then explored the role of enzymatic and structural Pdu signal sequences on MCP assembly by deleting their encoding sequences from the genome alone and in combination. Deleting enzymatic signal sequences decreased MCP formation, but this defect could be recovered in some cases by overexpressing genes encoding the knocked-out signal sequence fused to a heterologous protein. By contrast, deleting structural signal sequences caused similar defects to knocking out the genes encoding the full length PduM and PduB proteins. Our results contribute to a growing understanding of how MCPs form and function in bacteria and provide strategies to mitigate assembly disruption when encapsulating heterologous pathways in MCPs.

## Introduction

Biomanufacturing is a promising method to sustainably synthesize chemicals such as fuels, medicines, and materials. In contrast to traditional chemical synthesis, bioprocesses can operate at lower temperatures, use lower-value feedstocks, and access the wide range of molecules organisms have evolved to produce^1,2^. However, to achieve yields high enough to compete with traditional chemical production, metabolic engineering must overcome many challenges that limit pathway productivity, such as kinetic bottlenecks, toxicity of pathway products and intermediates to the host, and off-target product formation^3–6^. Spatial organization of enzymatic pathways is an attractive approach to address some of these challenges^7^. Successful strategies for enzyme colocalization have included employing synthetic DNA and protein scaffolds, which were used to increase the yields of L-threonine, mevalonate, 1,2- propanediol, and glucaric acid production pathways^8–11^. Microorganisms have also evolved an alternative spatial organization strategy known as bacterial microcompartments (MCPs), which are proteinaceous organelles used by many bacteria to encapsulate certain metabolic pathways. MCPs are roughly 100 to 200 nm in diameter and consist of a liquid-like enzymatic core encapsulated by a semipermeable polyhedral shell of self-assembling proteins^12,13^. MCPs include both carboxysomes, anabolic MCPs which encapsulate the RuBisCO enzyme required for CO_2_ fixation in cyanobacteria and chemoautotrophs, and metabolosomes, catabolic MCPs which encapsulate enzymatic pathways that metabolize niche carbon substrates^14^.

MCPs are a common spatial organization strategy in bacteria, as operons encoding metabolosomes have been identified in 45 of 83 bacterial phyla. The pathways encapsulated by metabolosomes metabolize a variety of substrates, but nearly all are hypothesized to pass through a toxic, volatile aldehyde intermediate^15^. By colocalizing pathway enzymes inside a diffusion barrier, metabolosomes are hypothesized to benefit the pathways they encapsulate by protecting the cell from toxic intermediates, increasing local intermediate concentrations to overcome slow enzyme kinetics, reducing competition with other cellular pathways, and providing private cofactor pools^16–20^. Engineering MCPs to encapsulate heterologous biosynthetic pathways has emerged as an attractive opportunity to impart the benefits of MCPs onto industrially relevant pathways, particularly those that share characteristics of natively encapsulated pathways such as intermediate toxicity, kinetic bottlenecks, and high cofactor requirements.

One of the best-studied metabolosomes is the 1,2-propanediol utilization (Pdu) MCP found in *Salmonella enterica* serovar Typhimurium LT2. This MCP encapsulates a pathway that converts 1,2-PD to propionate and 1-propanol through a toxic propionaldehyde intermediate (Figure 1a)^13^. Propionate can then be utilized by the cell’s central metabolism to produce energy in the form of ATP^21,22^. The proteins that form the Pdu MCP are expressed from the *pdu* operon in the *S. enterica* genome, which contains 22 genes (*pduA* through *pduX*) (Figure 1b)^13^. This operon expresses the self-assembling proteins that form the Pdu MCP shell^23^, pathway enzymes that convert 1,2-PD to propionate, cofactor recycling enzymes that regenerate adenosyl cobalamin (Ado-B_12_) and NADH within the MCP^24,25^, and several proteins of unknown function^12^.

**Fig 1.**
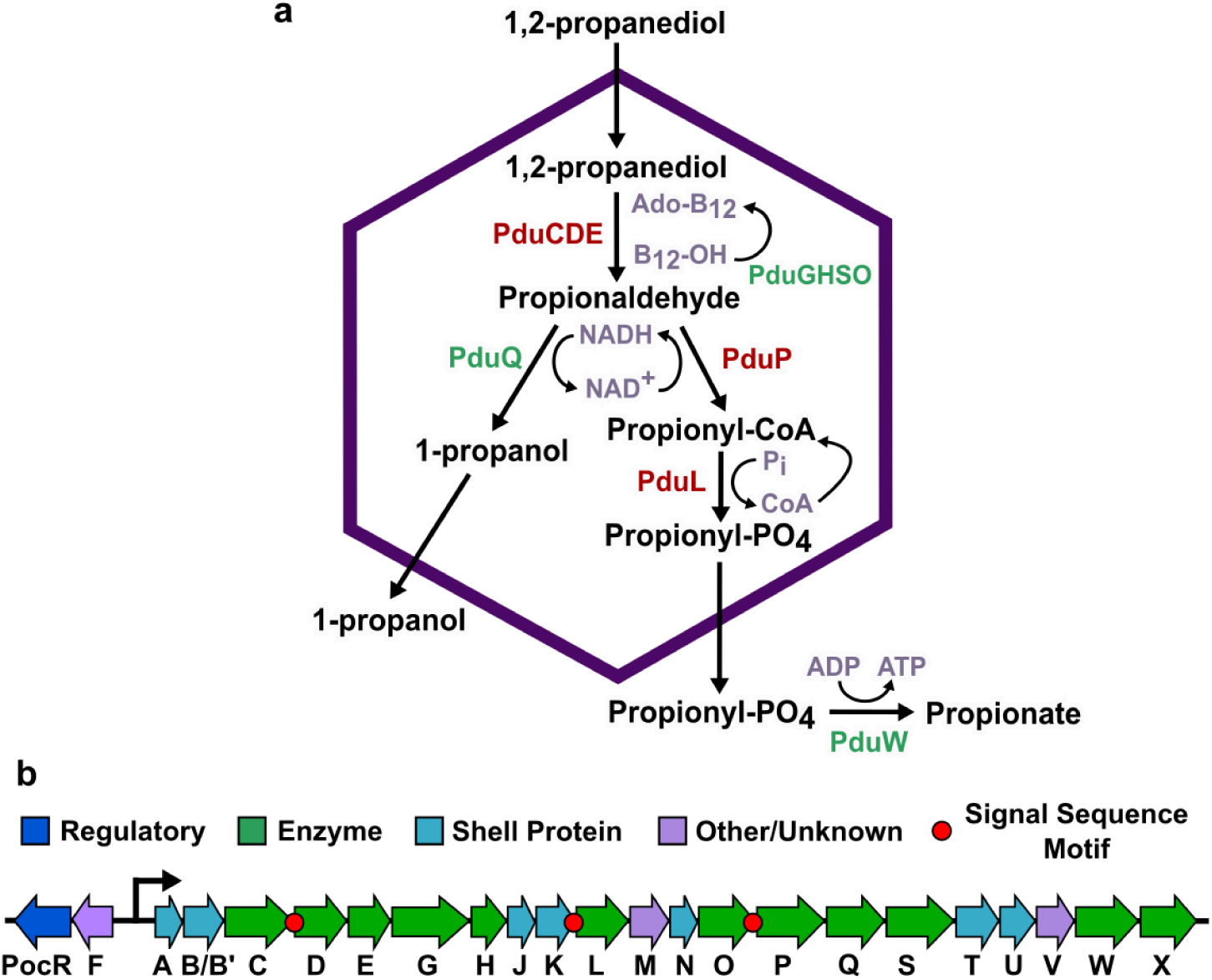
The 1,2-propanediol utilization microcompartment (Pdu MCP) pathway and operon. (a) The native Pdu MCP contains main pathway (red) and cofactor recycling (green) enzymes that degrade 1,2-propanediol to propionate and 1-propanol. (b) The *pdu* operon and adjacent genes in the *Salmonella enterica* genome encode the regulatory elements, structural proteins, and enzymes required to form the Pdu MCP, including three known signal sequences on the N-termini of the PduD, PduP, and PduL enzymes.

Several recent studies have used engineered self-assembling structures based on Pdu MCP shells as scaffolds or compartments to spatially organize and improve flux through several heterologous pathways^26,27^. Spatially organizing enzymes using these MCP-based structures has improved flux through several heterologous pathways, including pathways for 1,2-propanediol and ethanol production^28–30^. To minimize disruptions to MCP function when encapsulating heterologous pathways, it is critical to understand how manipulating different MCP components affects MCP assembly. Although the primary functions of most Pdu MCP components are known, how these functions unite to form functional MCP shells and cores is poorly understood. In particular, we do not understand how encapsulating heterologous cargo proteins in the MCP core might change MCP structure or interfere with proper MCP formation and function.

The exact mechanisms by which all MCP cargo are encapsulated are not well understood, but some MCP cargo proteins are known to contain encapsulation peptides, or signal sequences, that target them to the enzymatic core^31,32^. In addition to mediating encapsulation of native Pdu proteins, these signal sequences can also target heterologous proteins to the MCP lumen. Signal sequences appear to target the MCP core rather than any component of the shell, as they still colocalize with other components of the enzymatic MCP core when the core and shell are separated^33^. Many Pdu proteins without identified signal sequences also localize to the MCP core, but the encapsulation mechanisms for these proteins are unknown^25,33,34^. Three Pdu MCP enzymes contain characterized signal sequences, PduD^1–18^ (ssPduD), PduP^1–18^ (ssPduP), and PduL^1–20^ (ssPduL), which are necessary and sufficient for encapsulation of the PduCDE, PduP, and PduL enzymes (Fig. 1b). These signal sequences were identified by multiple sequence alignments of PduD, PduP, and PduL with homologues not associated with compartments. The Pdu-associated enzymes had N-terminal extensions that were not present in homologous enzymes, suggesting that the N-termini of these proteins may have structural roles related to their compartmentalization^31,35–37^.

Although the amino acid sequences of these signal sequences are poorly conserved (≤25% pairwise identity), they share a common motif of alternating pairs of hydrophobic and hydrophobic residues. This motif is widely conserved across MCP systems^32,38^, and several *de novo* signal sequences have been created based on this shared motif^39^. Previous studies have indicated that signal sequences fold into amphipathic α-helices^28,32,40,41^, and several studies have also suggested that they may mediate aggregation of the enzymes they are attached to^12,42^. While all three Pdu signal sequences share a highly conserved structure, they differ in encapsulation efficiency, which is the proportion of expressed signal sequence-tagged protein that is encapsulated in MCPs^43^.

In this study, we investigate the role of signal sequences in targeting proteins to the MCP core and how these encapsulation mechanisms influence MCP formation and structure. Because signal sequences are responsible for targeting many cargo proteins to the MCP core, we hypothesized that they may mediate interactions involved in MCP formation and the properties of the resulting MCPs. We first used amino acid sequence alignments and protein structure predictions to search the Pdu MCP for previously unidentified motifs resembling signal sequences. This search and subsequent characterizations revealed functional signal sequences on the structural MCP proteins PduM and PduB and showed that an N-terminal extension on PduE does not function as a signal sequence in this system. Because ssPduM and ssPduB are the first signal sequences discovered on structural proteins, we then knocked out the sequences encoding each of the Pdu signal sequences, alone and in combination, to characterize how the roles of enzymatic signal sequences in MCP formation differ from those of structural signal sequences. We found that removing enzymatic signal sequences, particularly ssPduD, decreased MCP formation, but this defect could be partially rescued by overexpressing the knocked-out signal sequence attached to GFP. This suggests that enzymatic signal sequences play roles in MCP assembly beyond just localizing enzymes to the MCP core. By contrast, removing structural signal sequences caused similar defects to full-length *pduM* and *pduB* knockouts, and these defects could not be rescued by overexpressing the knocked-out signal sequences. This suggests that these defects are caused by removing the bodies of PduM and PduB from the MCP, rather than by removing the signal sequences themselves. Finally, we mutated a region within two weak/nonfunctional Pdu signal sequences, ssPduL and ssPduE, to investigate whether such mutations can predictably increase signal sequence encapsulation efficiency. The results of our study provide additional tools for identifying putative MCP signal sequences based on protein structure rather than on comparisons with non-compartment associated homologues. In addition, because signal sequences are likely to be manipulated when encapsulating heterologous pathway enzymes in MCPs, our findings also provide design rules for how heterologous pathways can be encapsulated while minimizing disruptions to MCP formation.

## Results

### N-terminal extensions on PduM and PduB act as signal sequences, while an N-terminal extension on PduE does not

To comprehensively investigate the role of signal sequences in Pdu MCPs, we first set out to assemble a complete list of the Pdu signal sequences. To accomplish this, we searched Pdu proteins for any previously unidentified signal sequence motifs and performed more extensive testing on previously proposed signal sequence motifs. We first noticed that the N-terminus of PduM contains an amino acid motif similar to known Pdu signal sequences. PduM is a low-abundance structural protein that is highly conserved between *pdu* operons in different organisms but has no known sequence homology to any proteins outside of Pdu MCPs^12,44,45^. PduM localizes to the MCP core, and its absence causes partial separation of the MCP core and shell^12^. Because PduM lacks known sequence homologues outside of Pdu MCPs, we could not perform a sequence alignment of PduM with non-compartment associated homologues, the technique which was used to discover the signal sequences associated with PduD, PduL, and PduP^31,35,36,44^. Therefore, we instead used the HHpred program to search for structural homology between PduM and proteins with deposited structures in the Protein Data Bank^46^. HHpred detected structural homology between an N-terminal α-helix in PduM and N-terminal helices in PduD and PduE orthologs from *Klebsiella oxytoca* (PDB entry 1EEX, chains E and C)^47^. Because the PduD N-terminus contains a known signal sequence and the PduE N-terminus is known to form an amphipathic helix^36,40^, these HHpred hits suggest that the PduM N-terminus might contain a signal sequence as well. In addition, we noted that the PduM N-terminus contained alternating sets of hydrophobic and hydrophilic residues, similar to other signal sequences (Figure 2a). Interestingly, HHpred also detected structural homology between other portions of PduM and proteins with Rossmann-like folds, particularly flavin-binding proteins. Based on these results, we set out to determine if PduM contained a signal sequence necessary and sufficient for its encapsulation in Pdu MCPs.

**Fig 2.**
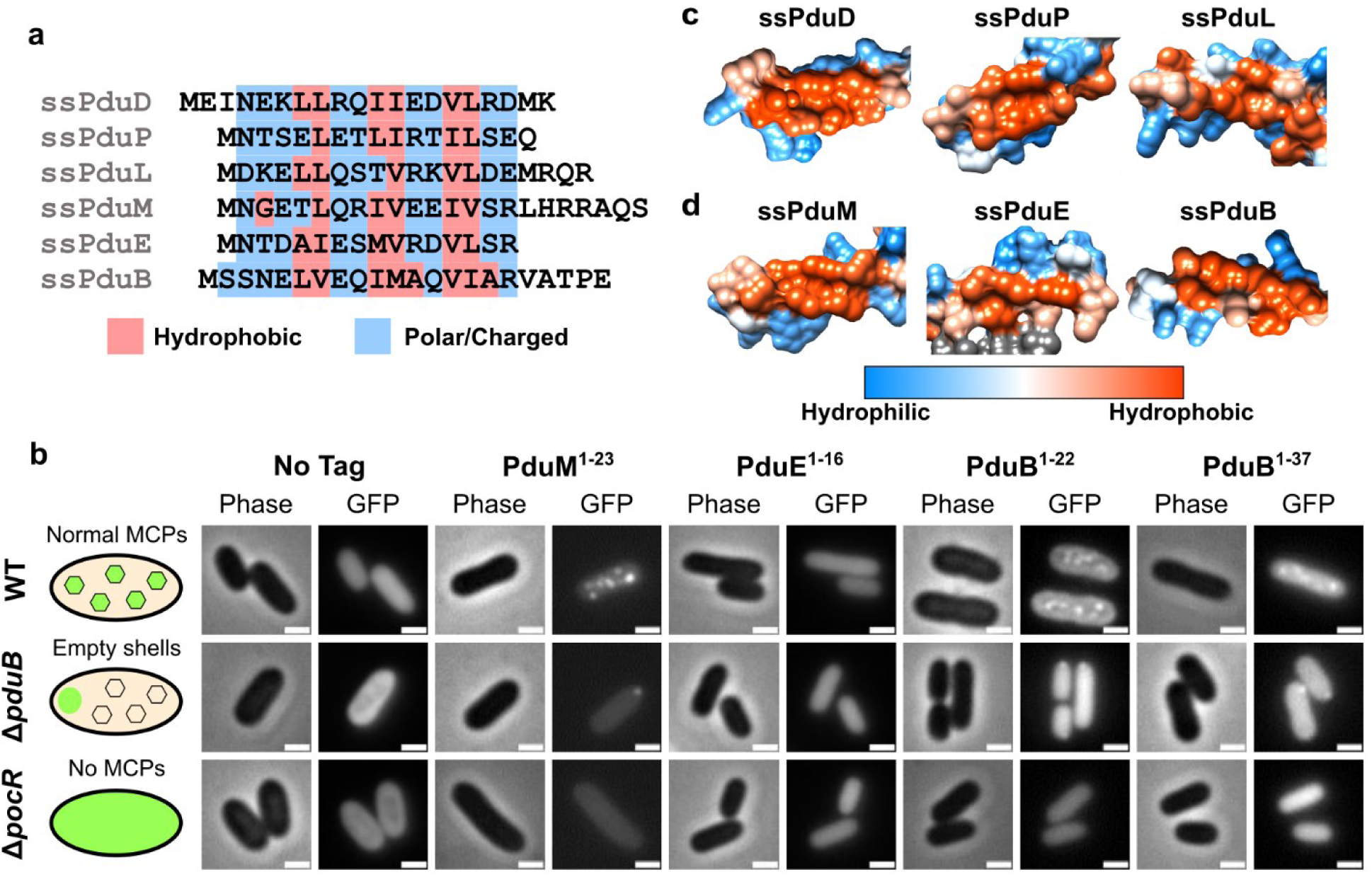
Characterization of N-terminal extensions on PduB, PduE, and PduM as encapsulation peptides. (a) Amino acid sequence alignment of the signal sequence-like motifs at the N-termini of PduD, PduP, PduL, PduM, PduE, and PduB. Hydrophobic residues are highlighted in red and hydrophilic residues are highlighted in blue, showing a pattern of alternating sets of hydrophobic and hydrophilic residues conserved between most sequences. (b) Predicted protein structures of previously identified signal sequences (ssPduD, ssPduP, and ssPduL) and (c) signal sequences characterized in this study (ssPduM, ssPduE, and ssPduB). These structures were downloaded from the AlphaFold Protein Structure Database and visualized using UCSF Chimera using Chimera’s default coloring for hydrophobicity of protein surfaces^51–53^. The hydrophobic sides of the amphipathic helices are shown. Hydrophilic areas are shown in blue and hydrophobic areas are shown in red. (d) Optical and fluorescence micrographs of putative signal sequences fused to GFP. These constructs were overexpressed both in MCP-forming *S. enterica* and in two assembly-deficient *S. enterica* strains (Δ*pduB* and Δ*pocR*). All scale bars are 1 μm. Similar results were observed across at least three biological replicates of each strain.

To test whether the N-terminus of PduM was sufficient to target cargo to the MCP core, we fused the putative PduM signal sequence (PduM^1–23^) to GFP and overexpressed this construct in *S. enterica* strains expressing both assembly competent and assembly deficient Pdu MCPs. Like other signal sequences, PduM^1–23^-GFP localized to fluorescent puncta in MCP-expressing wild-type *S. enterica*, suggesting that it associates with Pdu MCPs (Figure 2b). To determine whether PduM^1–23^-GFP was localizing to the MCP shell or core, we also expressed it in *S. enterica* lacking the shell protein PduB (Δ*pduB*), which causes decoupling of the MCP shell and core. In Δ*pduB*, the MCP core localizes to one of the cell poles while the shells form separately and are distributed throughout the cytoplasm^12,33,48^. Like other Pdu signal sequences, PduM^1–23^-GFP localized to polar bodies in Δ*pduB*, indicating that it associates with the MCP core rather than interacting directly with the shell. Finally, we expressed PduM^1–23^-GFP in a non-MCP-expressing strain that lacks PocR, the transcriptional activator of the *pdu* operon (Δ*pocR*) ^49,50^. We observed diffuse fluorescence when PduM^1–23^-GFP was expressed in Δ*pocR*, indicating that PduM^1–23^-GFP aggregation is dependent upon expression of other MCP components. Together, these results indicate that PduM^1–23^ is sufficient to target proteins to the Pdu MCP core.

We next investigated whether PduM^1–23^ is necessary for PduM encapsulation in Pdu MCPs. To do this, we fused both full-length PduM and PduM lacking its signal sequence motif (PduM^24-*^) to GFP and expressed these constructs in wild-type and Δ*pocR S. enterica*. In wild- type *S. enterica*, PduM-GFP overexpression resulted in multiple bright puncta per cell with low diffuse background (Supplementary Figure S1). Surprisingly, PduM^24-*^-GFP overexpression also occasionally gave rise to multiple puncta per cell, but these puncta were very dim with high diffuse background. Neither PduM-GFP or PduM^24-*^-GFP showed appreciable aggregation in Δ*pocR*. These results indicate that while PduM can still associate with MCPs to a low extent in the absence of its signal sequence, PduM^1–23^ is necessary to reach wild-type levels of PduM encapsulation.

Similarly to PduM, we observed that the N-terminus of PduE also contains alternating sets of hydrophobic and hydrophilic residues, and we therefore investigated its ability to act as a signal sequence (Figure 2a). This pattern in the PduE N-terminus was also previously noted by Kinney et al.^40^. A previous study found that like other Pdu enzymes that contain signal sequences, PduE contains an N-terminal extension that does not occur in non-compartment associated homologues, which suggests that this extension may play a compartmentalization-related role^37^. However, Fan and Bobik showed that this region is not responsible for localization of PduE to MCPs and rather is required for proper PduE enzymatic activity^36^. Although the N-terminal extension of PduE is not necessary for localizing PduE to MCPs, we hypothesized that it may still be sufficient to target heterologous proteins to MCPs outside of the context of PduE because it shares the same pattern of alternating hydrophobic and hydrophilic regions seen in other signal sequences. To test this hypothesis, we fused PduE^1–16^ to GFP and overexpressed this construct in *S. enterica* expressing Pdu MCPs. PduE^1–16^-GFP did not form fluorescent puncta in WT, Δ*pduB*, or Δ*pocR* strains, indicating that PduE^1–16^ does not act as a signal sequence in these contexts (Figure 2b).

Following the release of the AlphaFold Protein Structure Database, we were curious if the common motif shared by the Pdu signal sequences would be reflected by any similarities in their predicted structures. We therefore visualized the hydrophobicity surfaces predicted by AlphaFold for Pdu proteins containing signal sequence motifs. The predicted surfaces of PduD, PduP, PduL, and PduM showed that their signal sequences shared visibly similar structures, extending away from the body of the protein with similar hydrophobic surfaces on one side of the helix (Figure 2c, d)^51–53^. However, the predicted hydrophobicity surface of PduE showed that its N-terminal extension incorporates into the body of the protein instead of extending away from it (Supplementary Figure S2), consistent with findings that it is not a functional signal sequence and is instead required for proper enzymatic activity^36^.

Because the encapsulation mechanisms for many Pdu cargo proteins are still unknown, we searched the predicted structures of other proteins in the *pdu* operon for signal sequence-like structures. We found that the N-terminus of PduB, one of the MCP shell proteins, is predicted to fold into a structure resembling a signal sequence (Figure 2d). Although the PduB N-terminus does not follow the pattern of alternating sets of hydrophobic and hydrophilic amino acids as closely as the other Pdu signal sequences (Figure 2a), PduB^1–22^ is predicted to fold into an amphipathic helix with a hydrophobic pocket. PduB^23–37^ is predicted to form an unstructured linker between this helix and the body of the PduB protein, which interacts with the other MCP shell components. We therefore hypothesized that the N-terminus of PduB acts as a signal sequence and would be sufficient to target heterologous cargo to MCPs.

To test this, we fused PduB^1–22^ and PduB^1–37^ to GFP and overexpressed these constructs in *S. enterica* expressing Pdu MCPs. Like other signal sequences, PduB^1–22^-GFP and PduB^1–37^- GFP localized to fluorescent puncta in wild-type *S. enterica*, to polar bodies in Δ*pduB*, and were diffuse in Δ*pocR* (Figure 2b). However, the puncta formed by overexpression of these constructs were qualitatively dim compared to the diffuse background fluorescence, suggesting that PduB^1–22^ may interact with MCPs more weakly than other signal sequences. Previous studies have found that PduB^1–37^ deletions cause separation of the MCP core and shell^12,33,48^, indicating that PduB^1–37^ is necessary to link PduB, carrying with it the rest of the MCP shell, to the MCP core. In combination with our results, this suggests that PduB^1–37^ may bind the MCP shell to the core by the same mechanism other signal sequences use to bind cargo proteins to the MCP core. The characterization of ssPduM and ssPduB comprises the first report of encapsulation peptides on structural metabolosome proteins rather than enzymes. This result implies a broader view of signal sequences’ role in MCP assembly, beyond just encapsulation of cargo enzymes.

Finally, we assessed the encapsulation efficiencies of the signal sequence motifs on PduM, PduE, and PduB relative to known Pdu signal sequences. To accomplish this, we purified MCPs from strains overexpressing each signal sequence attached to GFP. We then assessed cargo encapsulation and expression by performing an anti-GFP western blot of the purified MCPs and whole cell lysates from these strains. Consistent with the ratios of punctate to diffuse fluorescence observed by microscopy, western blotting showed high encapsulation efficiencies for ssPduD-GFP, ssPduP-GFP, and ssPduM-GFP and much lower encapsulation efficiencies for ssPduL-GFP, ssPduB-GFP, and ssPduE-GFP (Supplementary Figure S3).

### Enzymatic signal sequences play essential and distinct roles in proper MCP formation

Identifying encapsulation peptides on structural proteins made us consider the role of signal sequences in overall MCP assembly and how these roles could differ between structural and enzymatic signal sequences. We also recognized that signal sequences are often manipulated when encapsulating heterologous cargo in MCPs, either by removing sequences encoding native signal sequences or by overexpressing signal sequences fused to heterologous proteins^30,43,54^. Therefore, understanding how signal sequences affect MCP assembly could advance efforts to engineer MCPs without disrupting their structure and function.

Because enzymatic signal sequences share a common structure suggested to contribute to the liquid-like properties of the MCP core, we hypothesized that removing these signal sequences might disrupt MCP assembly. To test this hypothesis, we combinatorially knocked out the sequences encoding the enzymatic signal sequences ssPduD, ssPduP, and ssPduL from the *S. enterica* genome and assessed the impact of these deletions on MCP formation using fluorescence microscopy and transmission electron microscopy (TEM) (Supplementary Figure S4a). We then overexpressed a suite of GFP reporters that localize to the MCP core in these strains. Previous studies have used fluorescence microscopy of encapsulated GFP reporters to show that MCP structural defects often change the number and spatial distribution of fluorescent puncta^33,55,56^. In wild-type *S. enterica*, core reporters typically form three to six fluorescent puncta distributed throughout the cytoplasm. When MCP assembly is disrupted such that shells either do not form or are separated from the core, the reporters typically localize to one or two fluorescent puncta located at the cell poles. If the reporters no longer localize to the MCP core at all, fluorescence is evenly distributed throughout the cytoplasm.

We first assessed how removing enzymatic signal sequences affects the number and spatial distribution of MCP cores by overexpressing GFP fused to the cofactor recycling enzymes PduG and PduO, which localize to the MCP core without a signal sequence, in the enzymatic signal sequence knockout strains^33^. Fluorescence microscopy of the PduG and PduO encapsulation reporters showed that the number of puncta per cell decreased in the enzymatic signal sequence knockout strains, but the spatial distribution of puncta was mostly unaffected (Figure 3a, b and Supplementary Figures S5, S6). All Δ*ssPduD* strains had an especially steep drop in puncta count, and, in general, the more signal sequences were knocked out, the larger the decrease in puncta count. PduO-GFP localized almost entirely to single polar bodies in Δ*ssPduD* strains, similar to its localization in Δ*pocR*, indicating that its encapsulation was particularly affected by the absence of ssPduD (Supplementary Figures S5, S6). These results suggest that the absence of enzymatic signal sequences decreases MCP core formation. In addition, because ssPduD has the highest encapsulation efficiency among the enzymatic signal sequences and targets the signature enzyme, PduCDE, to Pdu MCPs^36,43^, these results point to a possible correlation between a signal sequence’s encapsulation efficiency, significance in the MCP pathway, and importance in MCP assembly.

**Fig 3.**
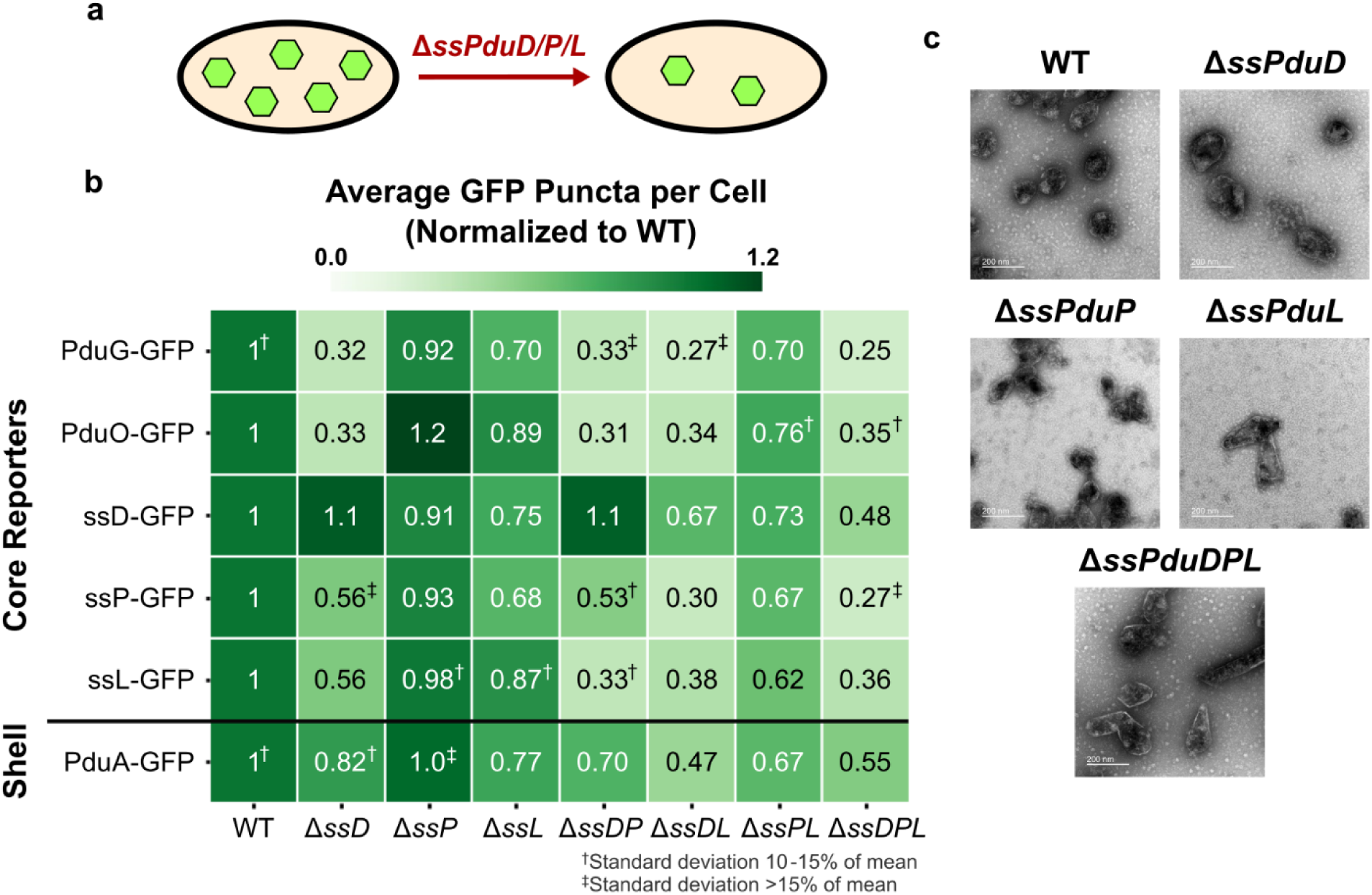
Impact of enzymatic signal sequences ssPduD, ssPduP, and ssPduL on MCP shell and core formation. (a) Knocking out the sequences encoding enzymatic signal sequences ssPduD, ssPduP, and ssPduL reduces formation of Pdu MCP shells and cores. (b) Average puncta observed per cell by fluorescence microscopy when Pdu signal sequences (ssD, ssP, ssL), cofactor recycling enzymes (PduG, PduO), and a shell protein (PduA) were fused to GFP and expressed in the enzymatic signal sequence knockout strains. Values shown in this figure were normalized by the average wild type puncta count for each reporter to facilitate comparison between reporters. Each value shown is the mean across three biological replicates, in which each replicate consisted of at least 30 cells counted from the same microscope slide. The standard deviation of the puncta count for each strain/reporter combination was less than 10% of the mean unless noted otherwise. Raw (i.e., not normalized) means and standard deviations of puncta counts are available in Supplementary Figure S6a. (c) Transmission electron micrographs of MCPs purified from wild type, Δ*ssPduD*, Δ*ssPduP*, Δ*ssPduL*, and Δ*ssPduDPL S. enterica*.

We also expressed PduA-GFP, a reporter that localizes to the MCP shell, in these strains to determine how removing enzymatic signal sequences affects MCP shell formation. PduA-GFP puncta counts also decreased as more enzymatic signal sequences were knocked out, following a pattern similar to the core reporters. These results indicate that knocking out signal sequences decreases overall MCP formation, rather than just formation of MCP cores.

Finally, we recognize that assembly defects observed when deleting signal sequences could result either directly from the absence of a signal sequence itself or because the body of its corresponding enzyme no longer localizes to MCPs. To distinguish between these possibilities, we investigated whether defects in enzymatic signal sequence knockout strains could be rescued by complementing one of the absent signal sequences fused to GFP, reintroducing the signal sequence while the body of its corresponding enzyme remained absent.

Interestingly, complementation with ssPduD-GFP resulted in higher puncta counts normalized to WT than all other encapsulation reporters in Δ*ssPduD* strains (ANOVA *p*<0.01, Supplementary Table S6). By contrast, ssPduD-GFP had similar normalized puncta counts to all but one strain/reporter combination in non-Δ*ssPduD* strains (ANOVA *p*>0.12, Supplementary Table S6). This suggests that overexpressing ssPduD-GFP can rescue assembly defects caused by knocking out ssPduD, but not defects due to knocking out other signal sequences. ssPduP- GFP and ssPduL-GFP complementation did not similarly recover puncta counts in Δ*ssPduP* and Δ*ssPduL* strains (Figure 3b). Normalized ssPduP-GFP and ssPduL-GFP puncta counts were significantly higher than other reporters in a few strains (ANOVA *p*<0.05, Supplementary Table S6) - for instance, ssPduP-GFP overexpression recovered puncta counts in Δ*ssPduD*Δ*ssPduP*, and ssPduL-GFP had a significantly higher normalized puncta count than ssPduP-GFP and PduG-GFP in Δ*ssPduD*Δ*ssPduL*. However, this pattern was not observed across most strains and reporters, so we conclude that ssPduP-GFP or ssPduL-GFP complementation does not recover puncta counts to as much of an extent as ssPduD-GFP. These results surprisingly suggest that different signal sequences may play different roles in supporting MCP assembly.

Because signal sequences share a common structure, we expected them to play similar roles in MCP assembly. If this were the case, overexpressing one signal sequence in multiple signal sequence knockout strains should rescue assembly equally across all strains. However, ssPduD- GFP recovers puncta counts only in Δ*ssPduD* strains, suggesting that the role of ssPduD in Pdu MCP assembly is unique from the roles of ssPduP and ssPduL.

We also performed TEM on purified MCPs from each of the knockout strains to more closely evaluate changes in MCP morphology (Figure 3c). All strains still formed shells and appeared to encapsulate cargo. MCPs from Δ*ssPduL* strains were slightly elongated, which may occur due to a polar effect that decreases the expression level of the downstream *pduN* gene^43^. Δ*ssPduD* and Δ*ssPduD*Δ*ssPduP* MCPs were qualitatively less homogeneous in shape than WT MCPs (Figure 3c, Supplementary Figure S7). No visible changes in MCP morphology were observed between WT and Δ*ssPduP* MCPs, which is consistent with the finding that Δ*ssPduP* puncta counts did not differ significantly from WT puncta counts. We also performed TEM of MCPs purified from Δ*ssPduD* and Δ*ssPduP* strains complemented with ssPduD-GFP and ssPduP-GFP to determine if recovery in puncta counts by fluorescence microscopy correlated with any changes in MCP morphology (Supplementary Figure S7). However, we did not notice any visible differences between MCPs with and without complementation, indicating that complementation rescues only the number of MCPs observed by optical microscopy and not the morphology of MCPs from these strains.

### Signal sequences are essential to the function of structural proteins PduM and PduB

We next knocked out the sequences encoding the structural signal sequences ssPduM and ssPduB to assess how the effects of removing structural signal sequences would differ from those of removing enzymatic signal sequences. Previous studies have shown that the MCP core and shell separate in strains lacking the PduB N-terminus^12,48^ because the body of the PduB protein (PduB^38-*^) can only bind to the shell^33^. Similarly, because PduM is a low-abundance structural protein, in contrast to the high-abundance enzymes encapsulated by ssPduD, ssPduP, and ssPduL, we expected that assembly defects in Δ*ssPduM* strains would occur because the body of PduM would be mostly unencapsulated (Supplementary Figure S1) rather than because of the absence of the signal sequence itself. Therefore, we hypothesized that knocking out the sequences encoding ssPduM and ssPduB would yield similar assembly defects to knocking out the full-length *pduM* and *pduB*, and these assembly defects would not be rescued by overexpressing the knocked-out signal sequences.

To test these hypotheses, we overexpressed ssPduD-GFP, PduG-GFP, ssPduM-GFP, and ssPduB-GFP in Δ*ssPduM*, Δ*pduM*, Δ*ssPduB*, Δ*pduB,* and Δ*ssPduM*Δ*ssPduB* (Δ*ssPduMB*) strains (Supplementary Figure S4b). ssPduD-GFP and PduG-GFP were included to report on MCP core formation. Because PduM and PduB play roles in proper connection of the MCP shell and core, we also included PduA-GFP as a shell protein reporter to assess how these knockouts impact shell formation. Puncta counts for ssPduB-GFP are not shown because its low encapsulation efficiency makes puncta dim and difficult to count (Supplementary Figure S8).

In Δ*ssPduM* and Δ*pduM*, all MCP core reporters localized mostly to polar bodies, with only a few puncta distributed throughout the cytoplasm (Figure 4a, Supplementary Figure S8). Puncta counts were similar across core reporters in these strains, indicating that overexpressing ssPduM-GFP could not recover assembly (Figure 4b). However, Δ*ssPduM* had lower puncta counts than Δ*pduM* (two-factor ANOVA p = 1.18×10^-5^), although this difference was only significant for some reporters (Supplementary Table S5). This suggests that when the body of PduM is present, but unencapsulated, it may still interact with the MCP by an unknown mechanism to cause a greater assembly defect than when PduM is completely absent. In Δ*pduB*, Δ*ssPduB*, and Δ*ssPduMB*, all MCP core reporters localized to polar bodies, indicating that assembly was disrupted and overexpressing ssPduB-GFP could not rescue proper MCP formation (Figure 4b, Supplementary Figure S8). ssPduM-GFP and ssPduB-GFP localize in a similar pattern as other encapsulation reporters that localize to the MCP core in all strains, which indicates that PduM and PduB are likely not required for each other’s encapsulation. This contradicts Yang et al.’s hypothesis that ssPduB and PduM bind to form a link between the MCP shell and core^12^, instead suggesting that ssPduB may directly target the shell to the core.

**Fig 4.**
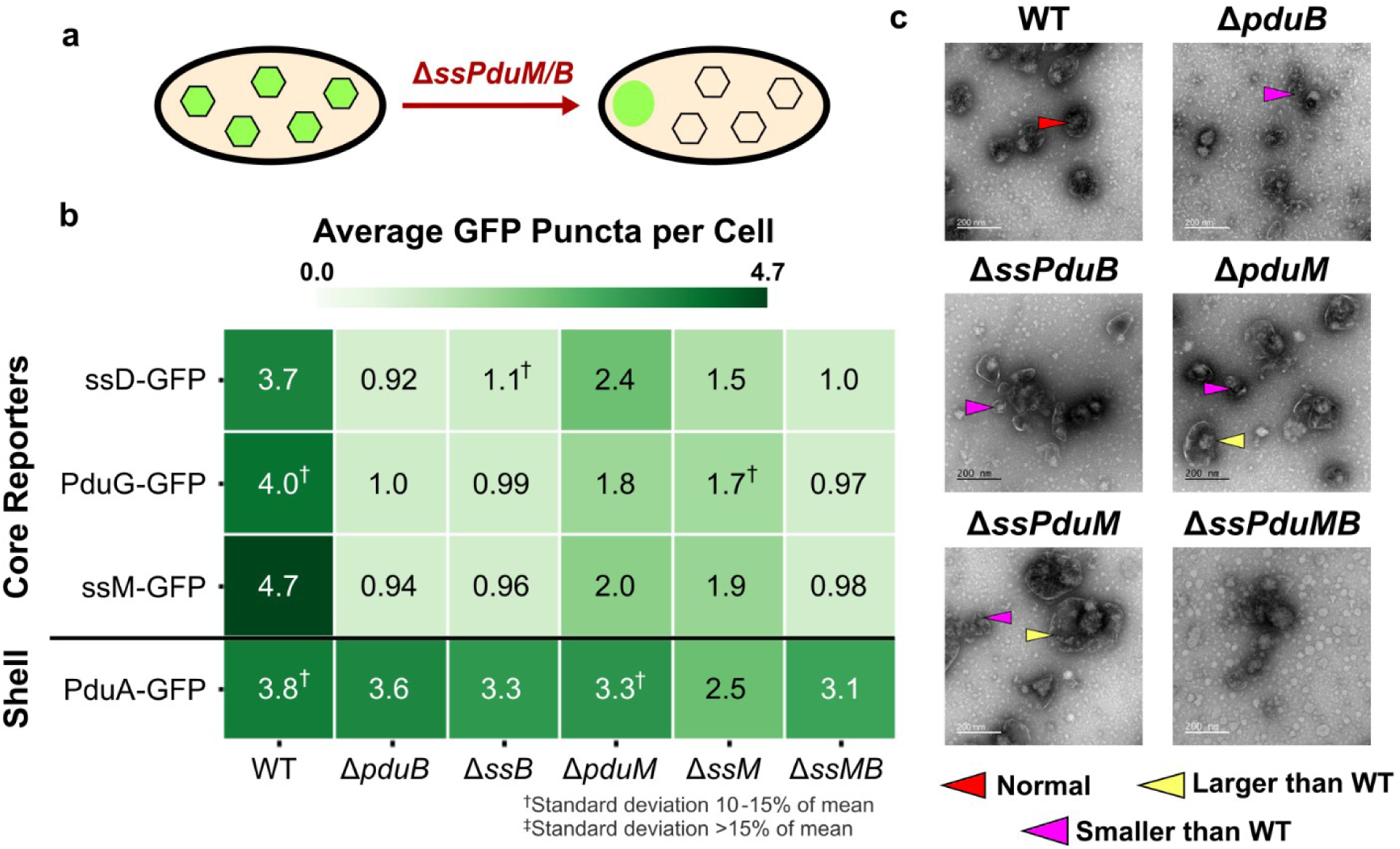
Roles of structural signal sequences ssPduM and ssPduB in MCP shell and core formation. (a) Knocking out sequences encoding the structural signal sequences ssPduM and ssPduB results in partial (ssPduM) or full (ssPduB) separation of Pdu MCP shells and cores. This causes core reporters to form an aggregate at one pole of the cell, while MCP shells remain distributed throughout the cytoplasm. (b) Average puncta observed per cell by fluorescence microscopy when Pdu signal sequences (ssD, ssM), cofactor recycling enzymes (PduG), and a shell protein (PduA) were fused to GFP and expressed in the structural signal sequence knockout strains. Each value shown in this figure is the mean across three biological replicates, in which each replicate consisted of at least 30 cells counted from the same microscope slide. The standard deviation of the puncta count for each strain/reporter combination was less than 10% of the mean unless noted otherwise. Means and standard deviations of puncta counts are available in Supplementary Figure S6b. (c) Transmission electron micrographs of MCPs purified from the structural signal sequence knockout strains. TEM imaging was performed on one biological replicate.

We next examined the impact of the structural signal sequences on shell formation by expressing the MCP shell reporter PduA-GFP in the structural signal sequence knockout strains. In strains where the MCP core and shell are connected, PduA-GFP should form similar numbers of puncta as encapsulation reporters. However, in strains where the core and shell are separated, PduA-GFP should form more puncta than the encapsulation reporters. When PduA-GFP was expressed in Δ*ssPduM* and Δ*pduM*, it formed significantly more puncta than the encapsulation reporters did (ANOVA *p*<0.01), which agrees with the partial separation of the MCP core and shell observed by Yang et al. in Δ*pduM*^12^. However, while the PduA-GFP puncta counts are similar to WT in Δ*pduM* and Δ*ssPduMB*, they are significantly lower than WT in Δ*ssPduM* (ANOVA p<0.001), indicating a reduced efficiency of shell formation in Δ*ssPduM*. PduA-GFP formed similar numbers of puncta when expressed in Δ*ssPduB*, Δ*pduB,* Δ*pduM*, Δ*ssPduMB*, and WT (ANOVA *p*>0.19). This result indicates that a proper number of MCP shells formed in these strains. Added to the result that core reporters localize to polar bodies in Δ*ssPduB* and Δ*pduB*, this suggests the shells were disconnected from the core in Δ*ssPduB* and Δ*pduB* (Figure 4b). These results suggest that when the shell and core are at least partially connected, MCP shell formation is disrupted by cytosolic PduM, but it is less disrupted when PduM is completely absent or when the core and shell are fully separated.

We performed TEM on purified MCPs from each of the knockout strains to more closely evaluate changes in MCP morphology (Figure 4c). Kennedy et al. reported that WT Pdu MCPs ranged from ∼60 to ∼170 nm in diameter when imaged by TEM, with an average diameter of approximately 100 nm^57^. Some Δ*ssPduM* and Δ*pduM* MCPs were well above this WT MCP size range, with some MCPs over 200 nm in diameter (Supplementary Figure S9). This suggests that PduM may play a role in regulating MCP size. Δ*pduB*, Δ*ssPduB*, and Δ*ssPduMB* had qualitatively lower electron density inside their MCPs, consistent with other results that indicate these strains form empty shells. Δ*ssPduMB* MCP shells in particular were less defined than MCPs from other strains. MCPs formed in Δ*pduB* and Δ*ssPduB* strains also generally appeared smaller than WT MCPs, consistent with previous studies showing that empty Δ*pduB* MCPs were smaller than WT MCPs and suggesting that the presence of cargo may influence MCP size^33^.

### Knocking out enzymatic signal sequences in combination with structural signal sequences reduces MCP shell formation

Next, we knocked out structural signal sequences in combination with the enzymatic signal sequences to assess whether this would cause additional assembly defects beyond knocking out structural signal sequences alone. Because Δ*ssPduB* and Δ*ssPduM* cause large structural defects that decouple the MCP core and shell, we hypothesized that this would outweigh structural defects caused by knocking out the enzymatic signal sequences, and knocking out enzymatic and structural signal sequences together would therefore not cause additional defects. However, because signal sequences are hypothesized to play a role in MCP core aggregation, we also hypothesized that knocking out all five signal sequences might affect the ability of signal sequences and other core enzymes to localize to the MCP core.

To test these hypotheses, we overexpressed fluorescent reporters for the MCP core (ssPduD-GFP and PduG-GFP) and shell (PduA-GFP) as well as the structural signal sequences (ssPduM-GFP and ssPduB-GFP) in Δ*ssPduD*Δ*ssPduP*Δ*ssPduL*Δ*ssPduM* (Δ*ssPduDPLM*), Δ*ssPduD*Δ*ssPduP*Δ*ssPduL*Δ*ssPduB* (Δ*ssPduDPLB*) and Δ*ssPduD*Δ*ssPduP* Δ*ssPduL*Δ*ssPduM*Δ*ssPduB* (Δ*ssPduDPLMB*) strains (Supplementary Figure S4b). In Δ*ssPduDPLM*, all MCP core reporters localized mostly to polar bodies, similar to their localization in Δ*ssPduM* and Δ*pduM* (Figure 5a, Supplementary Figure S8), suggesting that structural defects in the MCP core caused by knocking out ssPduM largely outweigh structural defects caused by knocking out the enzymatic signal sequences (Figure 5b). In Δ*ssPduDPLMB* and Δ*ssPduDPLB*, all core reporters localized to polar bodies, similar to their localization in Δ*ssPduB* and Δ*pduB* (Figure 5a, Supplementary Figure S8). This indicates that the structural defects in the MCP core caused by knocking out ssPduB outweighed defects caused by knocking out any other signal sequences. Interestingly, the presence of polar bodies in Δ*ssPduDPLMB* also shows that MCP cargo can still colocalize to an aggregate without any signal sequences. ssPduD-GFP puncta counts were not significantly different than other core reporters in Δ*ssPduDPLM*, Δ*ssPduDPLB*, or Δ*ssPduDPLMB* (ANOVA *p*>0.2), indicating that ssPduD-GFP overexpression did not recover puncta counts as it did in *ΔssPduD* strains not combined with structural signal sequence knockouts.

**Fig 5.**
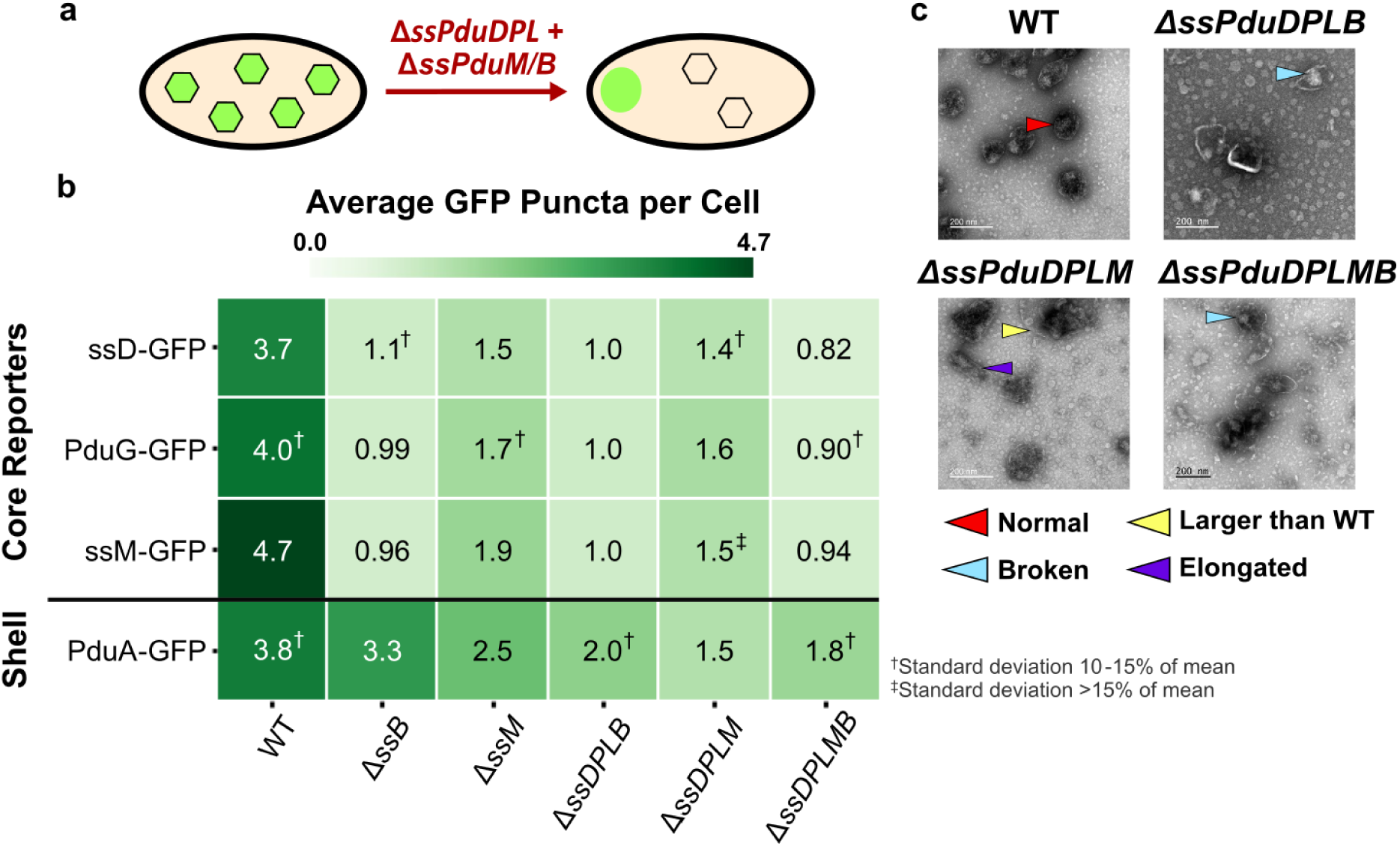
Impact of enzymatic signal sequence (ssPduD, ssPduP, ssPduL) deletions in combination with structural signal sequence (ssPduM, ssPduB) deletions on MCP shell and core formation. (a) Knocking out the sequences encoding enzymatic signal sequences ssPduD, ssPduP, and ssPduL in combination with those encoding the structural signal sequences ssPduM and ssPduB results in partial (ssPduM) or full (ssPduB) separation of Pdu MCP shells and cores and a decrease in MCP shell formation. Core reporters form an aggregate at one pole of the cell, while MCP shells remain distributed throughout the cytoplasm. (b) Average puncta observed per cell by fluorescence microscopy when Pdu signal sequences (ssD, ssM), cofactor recycling enzymes (PduG), and a shell protein (PduA) were fused to GFP and expressed in the enzymatic + structural signal sequence knockout strains. Each value shown in this figure is the mean across three biological replicates, in which each replicate consisted of at least 30 cells counted from the same microscope slide. The standard deviation of the puncta count for each strain/reporter combination was less than 10% of the mean unless noted otherwise. Means and standard deviations of puncta counts are available in Supplementary Figure S6b. (c) Transmission electron micrographs of MCPs purified from the enzymatic + structural signal sequence knockout strains. TEM imaging was performed on one biological replicate.

PduA-GFP formed significantly fewer puncta when expressed in Δ*ssPduDPLB*, Δ*ssPduDPLM*, and Δ*ssPduDPLMB* strains than in WT (ANOVA *p*<0.0001, Figure 5b). However, PduA-GFP still formed significantly more puncta than core reporters when expressed in Δ*ssPduDPLB* and Δ*ssPduDPLMB*, indicating separation of the shell and core (ANOVA *p*<0.001). Δ*ssPduDPLB*, Δ*ssPduDPLM*, and Δ*ssPduDPLMB* also had significantly lower PduA-GFP puncta counts than Δ*ssPduB*, Δ*ssPduM*, and Δ*ssPduMB*, respectively (ANOVA *p*<0.05). These results suggest that knocking out enzymatic and structural signal sequences together reduces MCP shell formation.

Finally, we performed TEM on purified MCPs from each of the knockout strains to more closely evaluate changes in MCP morphology (Figure 5c). Similar to Δ*ssPduM* and Δ*PduM* MCPs, Δ*ssPduDPLM* MCPs still had clearly defined shells and formed some MCPs >200 nm in diameter, above the reported size range of WT MCPs observed by TEM (Supplementary Figure S9)^57^. In contrast, we did not observe any fully closed, unbroken shells in Δ*ssPduDPLB* and Δ*ssPduDPLMB* MCPs across multiple biological replicates. In combination with the reduction in PduA-GFP puncta counts in these strains, this result suggests that removing enzymatic signal sequences in combination with separating the MCP shell and core may disrupt formation of complete MCP shells.

### Mutating signal sequences causes unpredictable changes in their encapsulation efficiencies

The Pdu signal sequences vary widely in encapsulation efficiency, but we could not find any immediately apparent elements of their amino acid sequences or predicted structures that correlated with their encapsulation efficiencies. We therefore sought to determine if Pdu signal sequences could be mutated to predictably alter their encapsulation efficiencies. From our amino acid alignment of the Pdu signal sequences, we chose to mutate a four amino acid region whose properties differed between strong and weak signal sequences (Figure 6a). In the strong signal sequences ssPduD, ssPduP, and ssPduM, these four amino acids comprise two polar or charged residues followed by two hydrophobic leucine, isoleucine, or valine residues. However, in the weak signal sequence ssPduL, the first hydrophobic residue is substituted by a polar threonine residue, and in the non-functional signal sequence ssPduE, one of the leucine/isoleucine/valine residues is replaced with methionine. This methionine (M9) is also one of the residues predicted to interact with the body of the PduE protein (Supplementary Figure S2). We therefore hypothesized that mutating this distinct sequence of four amino acids may predictably modulate the encapsulation efficiency of the Pdu signal sequences. ssPduB was excluded from this experiment because its amino acid sequence does not align well with the other Pdu signal sequences. To test this hypothesis, we designed six mutated signal sequences in which these four amino acids in ssPduE (ESMV) and ssPduL (QSTV) were replaced with the corresponding amino acids from ssPduD (RQII), ssPduP (ETLI), and ssPduM (QRIV) (Figure 6a). We fused those ssPduE and ssPduL variants to GFP and overexpressed these constructs in wild-type (normal MCP formation), Δ*pduB* (empty MCP shells), and Δ*pocR* (no MCP expression) *S. enterica*.

**Fig 6.**
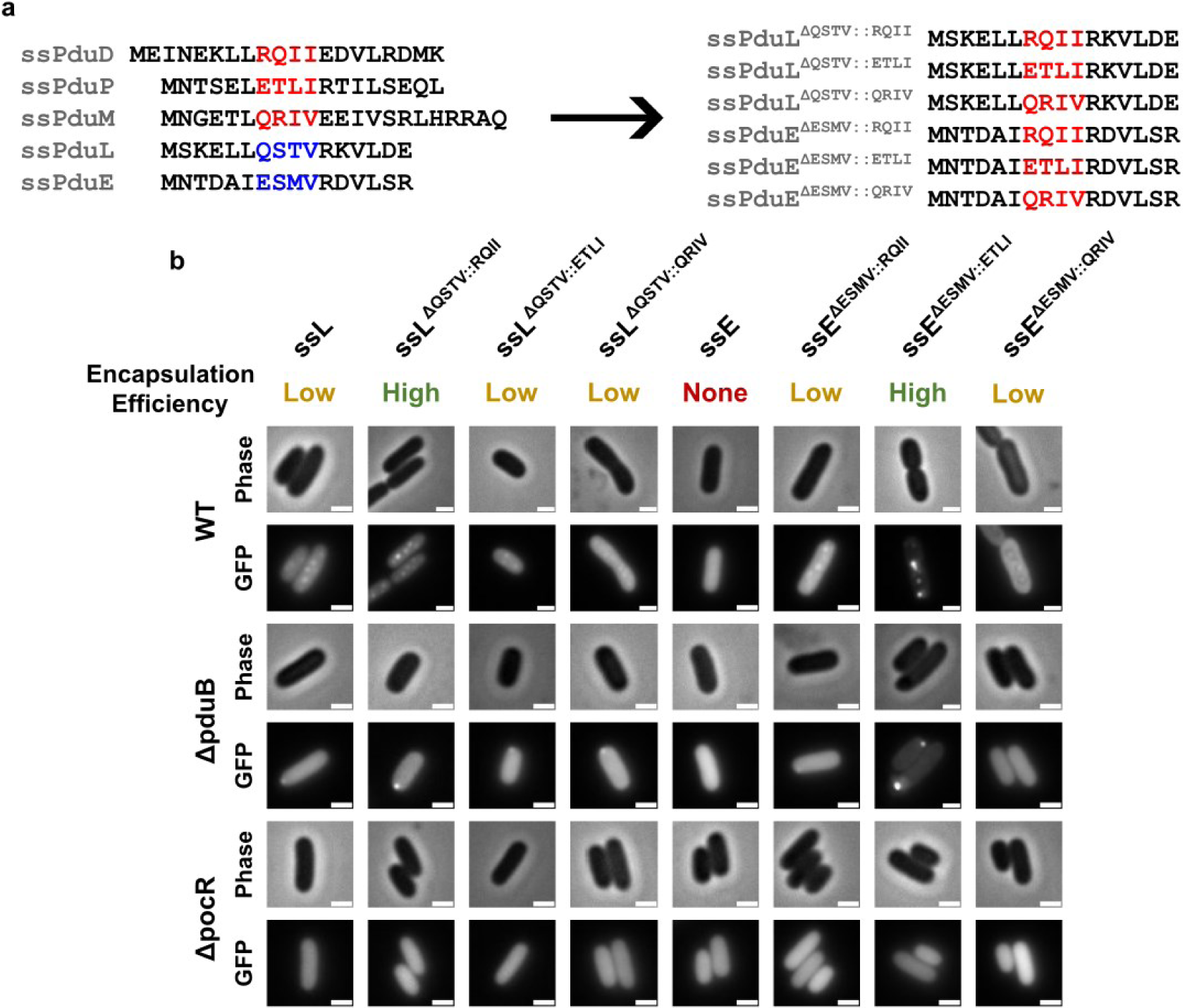
Mutations in a four amino acid region alter the encapsulation efficiencies of ssPduL and ssPduE. (a) Amino acid sequence alignment of the native (left) and mutated (right) Pdu signal sequences. Four amino acids in ssPduL and ssPduE were replaced with the corresponding residues from ssPduD, ssPduP, and ssPduL. (b) Optical and fluorescence micrographs of the ssPduL and ssPduE mutants fused to GFP. These constructs were overexpressed both in MCP-forming *S. enterica* and in two assembly-deficient *S. enterica* strains (Δ*pduB* and Δ*pocR*). All scale bars are 1 μm. Similar results were observed across at least three biological replicates.

We observed punctate fluorescence when overexpressing ssPduE^ΔESMV::RQII^-GFP and ssPduE^ΔESMV::ETLI^-GFP in WT *S. enterica*. Overexpressing ssPduE^ΔESMV::QRIV^-GFP also gave rise to noticeable, but extremely dim, punctate fluorescence (Figure 6b). These results confirm that ssPduE is “close” to a functional signal sequence, as substitution of just a small number of residues makes it capable of encapsulation. Overexpression of all ssPduL variants in WT *S. enterica* also resulted in puncta. Out of all ssPduE and ssPduL variants, ssPduE^ΔESMV::ETLI^-GFP and ssPduL^ΔQSRV::RQII^-GFP gave rise to particularly high punctate fluorescence with lower diffuse background, suggesting higher encapsulation efficiencies for these two constructs. Accordingly, we identified polar bodies when all variants that resulted in punctate fluorescence in WT were expressed in Δ*pduB*, confirming that these constructs aggregate with other MCP core proteins. We observed diffuse fluorescence in Δ*pocR* with all constructs, showing that they cannot aggregate without expression of other MCP proteins. Together, our results indicate that these four amino acids do affect the encapsulation efficiency of signal sequences, but not in a predictable manner. Mutating these four positions in the weak/nonfunctional signal sequences ssPduE and ssPduL to match the pattern seen in the strong signal sequences ssPduD, ssPduP, and ssPduM can increase their encapsulation efficiencies, but not all such mutations do increase encapsulation efficiency. Further, when the same set of mutations were made across different signal sequences, the effects of these mutations on encapsulation efficiency were not consistent. These results are not unexpected given the highly sensitive and unpredictable relationship between protein folding and function.

## Discussion

MCPs may be useful for metabolic engineering of heterologous pathways that share characteristics with natively encapsulated pathways, such as toxic intermediates, high cofactor requirements, and kinetic bottlenecks^7^. However, MCPs are highly complex, self-assembling protein structures that rely on many critical and interconnected interactions. Successfully engineering these structures to encapsulate heterologous pathways will therefore require an understanding of which modifications can be made without upsetting these interactions. Signal sequences that target proteins to the MCP core are particularly likely to be modified when encapsulating heterologous pathways, as they are often used to target heterologous proteins to the MCP lumen and may be knocked out if heterologous proteins are expressed genomically from the *pdu* operon^28,30,43^. In this study, we therefore examined the properties of Pdu signal sequences and the roles they play in MCP assembly.

While investigating the properties of the Pdu signal sequences, we identified two novel signal sequences on the MCP structural proteins PduB and PduM, and we demonstrated that despite aligning well with known signal sequences, an N-terminal extension on PduE is not capable of targeting heterologous cargo to MCPs. ssPduM and ssPduB are the first metabolosome signal sequences identified on structural proteins rather than encapsulated enzymes. In contrast to previously discovered metabolosome signal sequences^31,35,36^, ssPduM and ssPduB could not have been found through sequence alignments with non-compartment associated homologues as they were attached to uniquely microcompartment-associated structural proteins^44^. Instead, we predicted that ssPduM may behave as a signal sequence by analyzing its structural homology and by aligning its amino acid sequence with known Pdu signal sequences. We then predicted ssPduB to behave as a signal sequence by noting that its structure qualitatively aligned with known signal sequences even though its amino acid sequence did not. These results suggest that encapsulation peptide activity is a consequence of the way a protein folds rather than its amino acid sequence alone. Our results showing that mutations in ssPduL and ssPduE cause inconsistent and unpredictable changes in encapsulation efficiency reinforce the conclusion that sequence is not the sole determinant of encapsulation activity.

The presence of ssPduM and ssPduB expands our understanding of signal sequences’ roles in MCP assembly, indicating that signal sequences are responsible not only for targeting cargo enzymes to the MCP core but also play a second role in MCP assembly through proteins that link the MCP core and shell. Although many mechanisms for the connection of the MCP shell and core have been proposed, our work further elucidates the mechanism by which PduB links the shell and core. The body of PduB binds to other tiles in the MCP shell^33^, while the N-terminal signal sequence ssPduB binds to the core. This also strengthens evidence by Lehman et al. and Kennedy et al. that PduB is the main component responsible for connecting MCP the shell and core^33,48^.

The characterization of ssPduM, ssPduE, and ssPduB also demonstrate that analyzing predicted MCP protein structures may be a more comprehensive way to identify putative signal sequences than searching MCP proteins for extensions that do not occur in non-compartment associated homologues. Our investigation of the signal sequence-like motif ssPduE also suggests that analyzing predicted protein structures could more accurately identify putative signal sequences. Even though PduE contains an extension that does not occur in non-compartment associated homologues^36,37^, its predicted structure visibly differs from other signal sequences, consistent with the result that ssPduE is not a functional signal sequence. In addition, this work showcases how protein structure prediction programs such as AlphaFold can be applied to identify and predict protein structure/function relationships, a process that may be broadly useful for other families of proteins and in protein engineering applications.

Identifying signal sequences on structural MCP proteins also caused us to question if signal sequences themselves, not just the proteins they are attached to, could be required for proper MCP formation and function. To investigate the roles signal sequences play in MCP assembly, we knocked out enzymatic and structural signal sequences both alone and in combination. The number of MCPs formed per cell dropped as enzymatic signal sequences are knocked out, particularly in Δ*ssPduD* strains. However, overexpressing ssPduD attached to GFP partially recovered MCP formation in Δ*ssPduD* strains. This suggests that the decrease in MCP formation in Δ*ssPduD* strains likely occurs because of absence of the signal sequence itself, rather than because the body of the PduD enzyme no longer localizes to MCPs. It remains unclear why removing signal sequences causes a defect in MCP assembly and why this defect can only be recovered by overexpression of ssPduD-GFP, rather than other signal sequences, in Δ*ssPduD* strains.

Knocking out the sequences encoding ssPduM and ssPduB resulted in partial (ssPduM) or full (ssPduB) separation of the MCP shell and core, similar to defects observed in the full-length knockout strains Δ*pduM* and Δ*pduB*. Unlike the enzymatic signal sequence knockout strains, assembly defects in Δ*ssPduM* and Δ*ssPduB* strains could not be recovered by overexpressing the knocked-out signal sequences. This suggests that these defects are caused by disconnecting the bodies of the PduM and PduB proteins from the MCP core, rather than by removing the signal sequences themselves. In addition, Δ*ssPduM* reduced MCP shell formation more than the full-length knockout Δ*pduM*, suggesting that cytosolic PduM could still interfere with MCP assembly and adding to evidence that PduM has a unique and highly sensitive role in MCP formation. We also showed that the absence of PduM and ssPduM changes the size distribution of purified MCPs. Future studies are required to further investigate these phenomena and explore other potential functions and mechanisms of PduM. For example, more work is needed to confirm that these results are directly related to the role of PduM, rather than to polar effects with small impacts on expression from nearby loci such as *pduN*. Finally, knocking out enzymatic signal sequences in combination with structural signal sequences caused MCP assembly defects similar to those caused by knocking out structural signal sequences alone. This indicates that the assembly defects caused by knocking out structural signal sequences largely outweighed the effects of knocking out the enzymatic signal sequences. However, MCP shell formation did decrease in strains with both enzymatic and structural signal sequences knocked out compared to strains with only structural sequences knocked out, suggesting that enzymatic signal sequences may be required for proper MCP shell assembly.

By characterizing genomic knockouts of the enzymatic and structural signal sequences, our work also provides design rules that can be used to minimize disruptions to MCP assembly when encapsulating heterologous pathways in MCPs. For instance, our results indicate that ssPduD, and preferably all enzymatic signal sequences, should be present when encapsulating heterologous pathways in MCPs, either attached to native cargo or supplemented by overexpression. Structural signal sequences are also critical for proper MCP formation and should not be disrupted, unless intentionally to separate the MCP shell and core. The development of these rules therefore advances the field closer to successfully modifying MCPs for use in spatial organization and metabolic engineering of industrially relevant enzymatic pathways.

## Methods

### Plasmid Creation

All plasmids and primers are listed in Supplementary Tables S1 and S2. Plasmids containing ssPduD-GFP, ssPduP-GFP, ssPduL-GFP, PduG-GFP, and PduO-GFP were generated as previously described^33^. All other plasmids used in this study were generated by Golden Gate cloning^58^ using a pBAD33t parent vector (chloramphenicol resistance gene, p15A origin of replication, arabinose-inducible promoter)^55^. Each insert was amplified to add compatible BsaI cut sites flanking the gene of interest and purified using a PCR purification kit. The insert(s) and pBAD33t entry vector were then digested with Eco31I, ligated with T4 DNA ligase, and transformed into *E. coli* DH10b cells. The resulting clones were screened using green-white screening, and candidate plasmids were sequence verified using Sanger sequencing.

### Recombineering

All genomic edits were generated using λ Red recombineering^59^. All strains were first transformed with the pSIM6 plasmid, which contains the λ Red machinery and a carbenicillin (Cb) resistance marker^60^. pSIM6 is induced at 42°C to express λ Red machinery and is ejected from the cell at 37°C. The DNA inserts containing the desired genomic edits were either ordered from Twist Biosciences or amplified by overhang PCR (Supplementary Table S3). Each insert contained about 50 base pairs of homology to the *S. enterica* genome upstream and downstream of the desired insertion site. The pSIM6 plasmid was induced at 42°C, then cells were electroporated and transformed with the desired DNA insert. Strains were recovered either at 30°C to retain pSIM6 or at 37°C to remove it. Each genomic edit was created through two successive rounds of recombineering. In the first round, a *cat/sacB* cassette amplified from the TUC01 genome was inserted into the locus of interest. *cat* provides chloramphenicol (Cm) resistance, while *sacB* provides sucrose sensitivity^59^. After the first round of recombineering, cells were grown on lysogeny broth (LB) agar plates containing 10 μg/mL of Cm and 30 μg/mL of Cb to select for those that had successfully integrated *cat/sacB*. In the second round, *cat/sacB* was replaced with the desired insert. After this round, cells were grown on agar plates containing 6% (w/v) sucrose to select for those that had successfully removed *sacB*. Resulting colonies were streaked onto agar plates containing 10 μg/mL of Cm to confirm Cm sensitivity, then sequenced at the locus of interest to confirm insertion of the desired mutation.

### MCP Purification

MCPs were purified from *S. enterica* cultures using a differential centrifugation method adapted from Sinha et al.^44^. Briefly, overnight cultures were grown at 30°C, 225 rpm in 5 mL of lysogeny broth, Miller formulation (LB-M). Overnight cultures were then subcultured 1:1000 into NCE medium (29 mM potassium phosphate monobasic, 34 mM potassium phosphate dibasic, 17 mM sodium ammonium hydrogen phosphate) supplemented with 0.5% (w/v) succinate as a carbon source, 0.4% (v/v) 1,2-PD to induce MCP formation, 50 mM ferric citrate, and 1 mM magnesium sulfate. Subcultures were grown at 37°C, 225 rpm until they reached an OD_600_ between 1 and 1.5. For strains containing plasmids encoding MCP cargo, overnight cultures and subcultures were supplemented with chloramphenicol, and 0.02% (w/v) arabinose was added to subcultures after 14 to 16 hours of growth to induce cargo expression. Subcultures were induced for 5 hours before harvest.

After the target OD_600_ was reached, reserve samples were collected if needed, then the cultures were harvested at 5,000 × *g* for 5 min. Cell pellets were resuspended in 12.5 mL of lysis buffer (32 mM Tris-HCl, 200 mM potassium chloride (KCl), 5 mM magnesium chloride (MgCl_2_), 0.6% (v/v) 1,2-PD, 0.6% (w/v) n-Octyl-β-D-thioglucopyranoside (OTG), 5 mM β-mercaptoethanol, 0.8 mg/mL lysozyme, 0.04 units/mL DNase I, pH 7.5-8.0) and incubated at room temperature, 60 rpm for 30 min. Lysates were centrifuged twice at 12,000 × *g*, 4°C for 5 min to remove cell debris, and the resulting supernatant was spun at 21,000 × *g*, 4°C for 20 min in a swinging-bucket rotor to pellet the MCPs. The MCP pellets were washed with 5 mL of wash buffer (32 mM Tris-HCl, 200 mM KCl, 5 mM MgCl_2_, 0.6% (v/v) 1,2-PD, 0.6% (w/v) OTG, pH 7.5-8.0) and centrifuged again at 21,000 × *g*, 4°C for 20 min. The resulting pellets were resuspended in 150 μL of buffer B (50 mM Tris-HCl, 50 mM KCl, 5 mM MgCl_2_, 1% (v/v) 1,2-PD, pH 8.0) and centrifuged three times at 12,000 × *g*, 1 min to pellet any remaining cell debris. The concentrations of purified MCPs were then determined by a bicinchoninic acid assay (Thermo Scientific), and MCP samples were stored at 4°C until use.

### Transmission Electron Microscopy

Before sample deposition, 400-mesh copper grids with a Formvar/carbon film (EMS Cat# FCF400-Cu-50) were hydrophilized using a glow discharge cleaning system. 10 μL of purified MCP sample was then deposited onto each grid for 5 to 10 seconds and wicked away. Next, the samples were negative stained with 1% (w/v) aqueous uranyl acetate (UA) solution. 10 μL of UA was added to the grid and immediately wicked away. This was repeated once, then another 10 μL of UA was added to the grid for 4 minutes. The UA was then wicked away from the grid completely. After sample deposition, grids were imaged using a JEOL 1400 Flash transmission electron microscope with a Gatan OneView camera. Sizing of MCPs in TEM micrographs was performed using ImageJ as described by Kennedy et al^57^.

### Fluorescence Microscopy and Puncta Counting

Cultures for microscopy were prepared by first growing each strain at 37°C, 225 rpm in 5 mL of LB-M overnight. The overnight cultures were then subcultured 1:500 into 5 mL of LB-M media containing 0.4% (v/v) 1,2-PD to induce MCP formation. If strains contained fluorescent cargo expressed from a plasmid, overnight cultures and subcultures were supplemented with 34 μg/mL chloramphenicol, and arabinose was added to subcultures to induce expression of the plasmid. The arabinose concentration used for induction varied between reporter constructs. For most reporters, subcultures were induced with 0.02% (w/v) arabinose, but no arabinose (leaky expression only) was used to induce subcultures containing PduG-GFP, PduO-GFP, and PduA- GFP, as these reporters were prone to aggregation when expressed at high levels. After 6 hours of growth at 37°C, 225 rpm, the subcultures were prepared for microscopy. The cultures were concentrated 7:1, then placed onto Fisherbrand frosted microscope slides and covered with a 22 mm × 22 mm, No. 1.5 glass coverslip. Slides were imaged on a Nikon Eclipse Ni-U upright microscope using a 100x oil immersion objective, and images were captured with an Andor Clara digital camera. GFP fluorescence images were collected using a C-FL Endow GFP HYQ bandpass filter. Nikon NIS Elements software was used for image acquisition. All phase contrast images were collected using a 200 ms exposure time. Fluorescence images for most GFP reporters used a 100 ms exposure time, except ssPduL-GFP (200 ms), PduM-GFP (500 ms), PduM^24-*^-GFP and ssPduM-GFP (1 s), PduA-GFP (2 s), and PduG-GFP and PduO-GFP (3 s). Before puncta counting, image brightness and contrast were adjusted in ImageJ so that all puncta were clearly visible.

### SDS-PAGE and Western Blots

Protein samples were separated using sodium dodecyl sulfate polyacrylamide gel electrophoresis (SDS-PAGE). Purified MCP samples and reserved culture samples were diluted in Laemmli buffer and boiled at 95°C for 15 min. Purified MCPs were normalized by protein concentration (measured by bicinchoninic acid assay), and reserved culture samples were normalized by OD_600_ at harvest. Boiled MCP samples were loaded onto a 15% (w/w) polyacrylamide Tris-glycine gel such that 4 μg of protein was added to each well. The gel was run at 120 V for 90 min, then stained with Coomassie brilliant blue R-250. MCP samples were then re-normalized based on the intensities of their PduA and PduJ bands, then run on a second SDS-PAGE gel for western blotting.

For western blotting, normalized MCP and lysate samples were separated by SDS-PAGE, then transferred to a polyvinylidene fluoride (PVDF) membrane at 70 V for 20 min using a Bio-Rad Criterion Blotter. After transfer, the membrane was blocked using 5% (w/v) dry milk powder in tris-buffered saline with Tween-20 (TBS-T) buffer (20 mM Tris, 150 mM sodium chloride, 0.05% (v/v) Tween-20, pH 7.5) at room temperature for 1 hour. The membrane was incubated with an α-GFP primary antibody (Takara Bio cat# 632380, diluted 1:8000 in TBS-T) overnight at 4°C, then washed with TBS-T. A fluorescent goat anti-mouse IgG secondary antibody (LI-COR Biosciences cat# 926-68070, diluted 1:15000 in TBS-T) was applied to the membrane for 1 hour at room temperature. Finally, the membrane was washed with TBS-T and imaged using an Azure 600 imaging system.

### Statistical Analysis

All puncta counts shown are the means across three independent biological replicates for each strain/reporter combination. Each replicate consisted of at least 30 cells imaged from the same microscope slide. For puncta counting heatmaps in main figures, raw (i.e., not normalized) average per cell puncta counts and standard deviations are shown in the supplementary information.

Statistical analysis was performed using the Statistics and Machine Learning Toolbox in MATLAB R2024a. *α* = 0.05 was used as the threshold for statistical significance in all tests. One-factor ANOVAs were performed with Dunnett post-hoc tests if only pairwise comparisons to a control group were needed, and Bonferroni post-hoc tests were used if other pairwise comparisons were needed. Simple main effects for two-factor ANOVAs were calculated if the ANOVA returned a significant interaction *p*-value. Because the two-factor ANOVA only included two strains, the simple main effect for each reporter was calculated by conducting a two-tailed Student’s *t*-test between the puncta counts for that reporter in the two strains. All *F* and *t* statistics, *p* values, and degrees of freedom are shown in Supplementary Table S5. For all tests, the null hypothesis was that all strain/reporter combinations included in the test had the same average puncta count, and the alternative hypothesis was that at least one had a different average puncta count.

## Acknowledgements

We acknowledge and thank all members of the Tullman-Ercek lab for their helpful discussions and support regarding this paper. We thank Brett Palmero, Dr. Chris Jakobson, and Dr. Edward Kim for constructing several plasmids and strains used in this study, Lee Bristol for providing microscopy assistance, and Brett Palmero and Kris Rosentel for their help with data analysis. E.R.J. was funded in part by the Northwestern University Synthesizing Biology Across Scales program, a National Science Foundation National Research Traineeship program (NSF DGE-2021900). E.R.J. and D.T.E. were funded by the Northwestern University Materials Science and Engineering Center (NSF DMR-2308691). N.W.K. and T.M.N. were supported by a grant from the Department of Energy (DOE DE-SC0022180). C.E.M. was funded by a grant from the United States Army (W52P1J2193023). S.L. was funded by the Molecular Biophysics Training Program grant from the National Institutes of Health (NIH 5T32GM140995-04). S.C. was funded by the Northwestern University Synthetic Biology Research Experience for Undergraduates program, which is funded by the National Science Foundation (NSF DBI-2150269). G.A.R. was supported by the National Science Foundation Graduate Research Fellowship Program (NSF DGE-1842165). Molecular graphics and analyses performed with UCSF Chimera, developed by the Resource for Biocomputing, Visualization, and Informatics at the University of California, San Francisco, with support from the National Institutes of Health (NIH P41-GM103311).

## Author Contributions

E.R.J, C.E.M., N.W.K., T.M.N., G.A.R, and D.T.E. conceived this project. N.W.K. and T.M.N. performed initial proof-of-concept experiments, and C.E.M. generated many strains and plasmids used in this study. E.R.J, N.W.K., S.L., and S.C. performed experiments that generated data shown in this manuscript. E.R.J. and S.L. wrote the manuscript. E.R.J., N.W.K., C.E.M., S.L., S.C., T.M.N., and D.T.E. analyzed and interpreted experimental data. All authors reviewed and contributed to the manuscript.

## Materials & Correspondence

Correspondence and requests for materials should be addressed to Danielle Tullman-Ercek.

