## Supplemental Tables and Figures for "Signal sequences target enzymes and structural proteins to bacterial microcompartments and are critical for microcompartment formation"

Johnson *et al.*

**Supplementary Table S1.** Strains used in this study.

| Strain Number | Organism | Genotype | Abbreviation |
|---|---|---|---|
| DTE003 | *S. enterica* serovar Typhimurium LT2 | Wild type | WT |
| CEMS344 | *S. enterica* serovar Typhimurium LT2 | Δ*pduD*::*pduD*$^{19-*}$ | Δ*ssPduD* |
| CEMS361 | *S. enterica* serovar Typhimurium LT2 | Δ*pduP*::M-*pduP*$^{18-*}$ | Δ*ssPduP* |
| CEMS362 | *S. enterica* serovar Typhimurium LT2 | Δ*pduL*::*pduL*$^{17-*}$ | Δ*ssPduL* |
| CEMS351 | *S. enterica* serovar Typhimurium LT2 | Δ*pduD*::*pduD*$^{19-*}$ Δ*pduP*::*M-pduP*$^{18-*}$ | Δ*ssPduDP* |
| ERJ004 | *S. enterica* serovar Typhimurium LT2 | Δ*pduD*::*pduD*$^{19-*}$ Δ*pduL*::*pduL*$^{17-*}$ | Δ*ssPduDL* |
| ERJ005 | *S. enterica* serovar Typhimurium LT2 | Δ*pduP*::M-*pduP*$^{18-*}$ Δ*pduL*::*pduL*$^{17-*}$ | Δ*ssPduPL* |
| CEMS360 | *S. enterica* serovar Typhimurium LT2 | Δ*pduD*::*pduD*$^{19-*}$ Δ*pduP*::M-*pduP*$^{18-*}$ Δ*pduL*::*pduL*$^{17-*}$ | Δ*ssPduDPL* |
| CMJS256 | *S. enterica* serovar Typhimurium LT2 | Δ*pocR* | N/A |
| CEMS179 | *S. enterica* serovar Typhimurium LT2 | Δ*pduB* | N/A |
| CEMS342 | *S. enterica* serovar Typhimurium LT2 | Δ*pduM* | N/A |
| CEMS375 | *S. enterica* serovar Typhimurium LT2 | Δ*pduB*::*pduB*$^{\Delta 3-30}$ | Δ*ssPduB* |
| ERJ035 | *S. enterica* serovar Typhimurium LT2 | Δ*pduM*::M-*pduM*$^{23-*}$ | Δ*ssPduM* |
| ERJ198 | *S. enterica* serovar Typhimurium LT2 | Δ*pduM*::M-*pduM*$^{23-*}$ Δ*pduB*::*pduB*$^{\Delta 3-30}$ | Δ*ssPduMB* |
| ERJ116 | *S. enterica* serovar Typhimurium LT2 | Δ*pduD*::*pduD*$^{19-*}$ Δ*pduP*::M-*pduP*$^{18-*}$ Δ*pduL*::*pduL*$^{17-*}$ Δ*pduB*::*pduB*$^{\Delta 3-30}$ | Δ*ssPduDPLB* |
| ERJ039 | *S. enterica* serovar Typhimurium LT2 | Δ*pduD*::*pduD*$^{19-*}$ Δ*pduP*::M-*pduP*$^{18-*}$ Δ*pduL*::*pduL*$^{17-*}$ Δ*pduM*:: M-*pduM*$^{23-*}$ | Δ*ssPduDPLM* |
| ERJ117 | *S. enterica* serovar Typhimurium LT2 | Δ*pduD*::*pduD*$^{19-*}$ Δ*pduP*::M-*pduP*$^{18-*}$ Δ*pduL*::*pduL*$^{17-*}$ Δ*pduM*:: M-*pduM*$^{23-*}$ Δ*pduB*::*pduB*$^{\Delta 3-30}$ | Δ*ssPduDPLMB* |
| TUC01 | *E. coli* W3110 | gal490 pglΔ8 λcI857 Δ(cro-bioA) int<>cat/sacB | N/A |

**Supplementary Table S2.** Plasmids used in this study.

| Plasmid number | Description | Origin | Resistance |
|---|---|---|---|
| CMJ069 | pBAD33t-PduD$^{1-20}$ (ssPduD)-SR-GFPmut2 | p15A | Chloramphenicol |
| EYK208 | pBAD33t-PduP$^{1-17}$ (ssPduP)-SR-GFPmut2 | p15A | Chloramphenicol |
| NWKp041 | pBAD33t-PduL$^{1-20}$ (ssPduL)-SR-GFPmut2 | p15A | Chloramphenicol |
| EYK193 | pBAD-PduG-SR-GFPmut2 | p15A | Chloramphenicol |
| NWKp043 | pBAD33t-PduO-GS-GFP | p15A | Chloramphenicol |
| pBJP017 | pBAD33t-PduM$^{1-23}$ (ssPduM)-GFPmut2 | p15A | Chloramphenicol |
| pERJ011 | pBAD33t-PduB$^{1-37}$ (ssPduB with linker)-GS-GFPmut2 | p15A | Chloramphenicol |
| NWKp048 | pBAD33t-PduA-GS-GFPmut2 | p15A | Chloramphenicol |
| pCEM100 | pBAD33t-PduE$^{1-16}$ (ssPduE)-GS-GFPmut2 | p15A | Chloramphenicol |
| pERJ025 | GFPmut2 | p15A | Chloramphenicol |
| pERJ012 | pBAD33t-PduB$^{1-22}$ (ssPduB without linker)-GS-GFPmut2 | p15A | Chloramphenicol |
| pERJ014 | pBAD33t-ssPduE$^{\Delta ESMV::RQII}$-GS-GFPmut2 | p15A | Chloramphenicol |
| pERJ015 | pBAD33t-ssPduE$^{\Delta ESMV::QRIV}$-GS-GFPmut2 | p15A | Chloramphenicol |
| pERJ016 | pBAD33t-ssPduE$^{\Delta ESMV::ETLI}$-GS-GFPmut2 | p15A | Chloramphenicol |
| pERJ017 | pBAD33t-ssPduL$^{\Delta QSTV::ETLI}$-GS-GFPmut2 | p15A | Chloramphenicol |
| pERJ018 | pBAD33t-ssPduL$^{\Delta QSTV::RQII}$-GS-GFPmut2 | p15A | Chloramphenicol |
| pERJ019 | pBAD33t-ssPduL$^{\Delta QSTV::QRIV}$-GS-GFPmut2 | p15A | Chloramphenicol |
| pCEM119 | pBAD33t-PduM-GS-GFPmut2 | p15A | Chloramphenicol |
| pCEM120 | pBAD33t-M-PduM$^{24-*}$-GS-GFPmut2 | p15A | Chloramphenicol |
| pSIM6 | λ Red system repressed by cI857 | pSC101 *repA$^{ts}$* | Ampicillin |

**Supplementary Table S3.** Primers used in this study.

| Name | Purpose | Description | Sequence (5' → 3') |
|---|---|---|---|
| TMDP021 | Sequencing | Amplify from upstream of *pduD* locus (Fwd) | ggcaacaggttatcgcctgc |
| TMDP022 | Sequencing | Amplify from upstream of *pduD* locus (Rev) | ccctgcaggctgttcatgc |
| TMDP072 | Sequencing, recombineering | Amplify from upstream of *pduD* locus (Fwd) | catccagaaagccaagctaacc |
| TMDP073 | Sequencing, recombineering | Amplify from upstream of *pduD* locus (Rev) | cgtccagcgttttattggtgg |
| TMDP086 | Sequencing | Amplify from upstream of *pduP* locus (Fwd) | tcatttacagggaaaagtggtcacc |
| TMDP087 | Sequencing | Amplify from upstream of *pduP* locus (Rev) | tgcgccagaaagccatcg |
| TMDP088 | Sequencing | Amplify from upstream of *pduP* locus (Fwd) | tgtgacctggcggatgc |
| TMDP089 | Sequencing | Amplify from upstream of *pduP* locus (Rev) | tgcagagcctgcatttgc |
| TMDP043 | Sequencing | Amplify from upstream of *pduL* locus (Fwd) | tcagctgcaatctgtgtctgg |
| TMDP044 | Sequencing | Amplify from upstream of *pduL* locus (Rev) | cagaacagtgctggacagtcg |
| TMDP081 | Sequencing | Amplify from upstream of *pduL* locus (Fwd) | ctccgtcattgaacctgagc |
| TMDP082 | Sequencing | Amplify from upstream of *pduL* locus (Rev) | tgagctgtacgcggatgc |
| oCEM187 | Sequencing | Amplify from upstream of *pduM* locus (Fwd) | cgcagcggcatatccatatgtc |
| oCEM188 | Sequencing | Amplify from upstream of *pduM* locus (Rev) | cggtaaacagtgatgtggtgcc |
| oCEM189 | Sequencing | Amplify from upstream of *pduM* locus (Fwd) | gatcgcgggctgattttcaaca |
| oCEM190 | Sequencing | Amplify from upstream of *pduM* locus (Rev) | gatttttgcgtggagacaaccg |
| oMPV029 | Sequencing | Amplify from upstream of *pduB* locus (Fwd) | gtgggtgaagtgaaagccgta |
| oMPV030 | Sequencing | Amplify from upstream of *pduB* locus (Rev) | tccatcgccatcacttcttcg |
| oERJ006 | Sequencing, recombineering | Amplify from upstream of *pduB* locus (Fwd) | GAAAGCCGTACACGTCA TCCC |
| oERJ007 | Sequencing, recombineering | Amplify from upstream of *pduB* locus (Rev) | CCTGATTCACAGGGCGT TTCG |
| oCEM159 | Recombineering | Amplify *cat/sacB* with homology upstream of *pduM* (Fwd) | gccgctggtgccgataacccgcat gcctttgcccggctggtaggcccgc gTGTGACGGAAGAT |
| oCEM160 | Recombineering | Amplify *cat/sacB* with homology downstream of *pduM* (Rev) | gatttttgcgtggagacaaccgcgc ccgtgactcgtgccagatgcatgat ATCAAAGGGAAAA |

| | | | |
|---|---|---|---|
| oCEM171 | Recombineering | *pduM* knockout (Fwd) | ctggtgccgataacccgcatgcctttgcccggctggtaggcccgcgatgaaatattcaattaattaagcaggagtaaatcatgcatctggcacgagtcac |
| oCEM172 | Recombineering | *pduM* knockout (Rev) | gtgactcgtgccagatgcatgatttactcctgcttaattaattgaatatttcatcgcgggcct |
| TMDP003 | Recombineering | Amplify *cat/sacB* with homology upstream of *pduP* (Fwd) | CAGCGTTGAGCAGGACATGG |
| TMDP004 | Recombineering | Amplify *cat/sacB* with homology downstream of *pduP* (Rev) | CCAGATGTGCTTATTGGTAAAGCGC |
| TMDP023 | Recombineering | Amplify *cat/sacB* with homology upstream of *pduL* (Fwd) | aaaatgtccgcgccagaagg |
| TMDP024 | Recombineering | Amplify *cat/sacB* with homology downstream of *pduL* (Rev) | cgaagttgcgtaacgctcagc |
| oCEM161 | Recombineering | Amplify $pduD^{19-*}$ with homology upstream of *pduD* (Fwd) | aaacatccctggcgctcttgatcccaacgagattgattaaggggtgagaaatgaagggcagcgataaacc |
| oCEM162 | Recombineering | Amplify $pduD^{19-*}$ with homology downstream of *pduD* (Rev) | gtcccggaccatcgattcaattgcgtcggtattcatggagttatcctttatcaaagcgccacgcg |
| oCEM163 | Recombineering | Amplify $pduP^{18-*}$ with homology upstream of *pduP* (Fwd) | atggacatagcacagaccgccatcgcggctattaacgtgggaactcatcaatgaataccacgccggcg |
| oCEM164 | Recombineering | Amplify $pduP^{18-*}$ with homology downstream of *pduP* (Rev) | accgctgtacaaccgcgtttgtagtgagaaggtattcatcgcgacctcagttagcgaatagaaaagccgttgg |
| oCEM165 | Recombineering | Amplify $pduL^{17-*}$ with homology upstream of *pduL* (Fwd) | ggcgaacctcgtacgctttgcattcattccggcaagcgaggtgaagcgtaatgcgccagcgg |
| oCEM166 | Recombineering | Amplify $pduL^{17-*}$ with homology downstream of *pduL* (Rev) | atgcagccgggagacaatctcctcgacaatgcgctgcagggtttcgccgttcatcgcgggcct |
| oCEM195 | Recombineering | Amplify $pduB^{\Delta 3-30}$ with homology upstream of *pduB* (Fwd) | gccctcacaccgatgtagaaaaaatcttaccgaagggaattagccaatgagccaacctatacgagagacggctatgg |
| NWKo512 | Recombineering | Amplify $pduB^{\Delta 3-30}$ with homology downstream of *pduB* (Rev) | ttcgccagtgcttcaaatcttttcgatctcatgaatcagcctcgtgggtAtcagatgtaggacggacgatcgtttttcg |
| oERJ001 | Golden Gate cloning | Amplify $pduB^{1-37}$ and $pduB^{1-22}$ to add Golden Gate overhang (For) | AGTTACGGTCTCacatgagcagcaatgagctggtgg |
| oERJ002 | Golden Gate cloning | Amplify $pduB^{1-37}$ to add Golden Gate overhang | GTCAACGGTCTCAaaccagccgtctctcgtataggttggg |

| | | | |
|---|---|---|---|
| | | compatible with GS linker (Rev) | |
| oERJ003 | Golden Gate cloning | Amplify *pduB*$^{1-22}$ to add Golden Gate overhang compatible with GS linker (Rev) | GTCAACGGTCTCAaaccttccggcgttgccacacg |
| oBJP094 | Golden Gate cloning | Amplify GFPmut2 to add Golden Gate overhang compatible with *pduM*$^{1-23}$ (*ssPduM*) (Fwd) | AttGGTCTCAgagcAgtaaaggagaagaacttttcactgga |
| oBJP095 | Golden Gate cloning | Amplify oBJP097 to add Golden Gate overhang (Fwd) | attaggtctcacatgaacggcgaaaccctgcag |
| oBJP096 | Golden Gate cloning | Amplify oBJP097 to add Golden Gate overhang (Rev) | attaggtctcagctctgggcacggcgatg |
| oBJP097 | Golden Gate cloning | Oligo encoding *pduM*$^{1-23}$ (*ssPduM*) | atgaacggcgaaaccctgcagcgcattgtcgaggagattgtctcccggctgcatcgccgtgcccagagc |
| oBJP137 | Golden Gate cloning | Amplify GFPmut2 to add Golden Gate overhang and GS linker (Fwd) | AttaGGTCTCAggttctAgtaaaggagaagaacttttcactgg |
| NWKo569 | Golden Gate cloning | Amplify *pduA* to add Golden Gate overhang (Fwd) | AttGGTCTCACATGCAACAAGAAGCACTAGG |
| NWKo570 | Golden Gate cloning | Amplify *pduA* to add Golden Gate overhang and GS linker (Rev) | ATTGGTCTCAGCTGCCttggctaattcccttcgg |
| NWKo506 | Golden Gate cloning | Amplify GFPmut2 to add Golden Gate overhang compatible with GS linker (Fwd) | AttGGTCTCACAGCAgtaaaggagaagaacttttcactgg |
| NWKo507 | Golden Gate cloning | Amplify GFPmut2 to add Golden Gate overhang (Rev) | ATTGGTCTCATTTAtttgtatagttcatccatgccatgtg |
| oCEM207 | Golden Gate cloning | Oligo containing *pduE*$^{1-16}$ (*ssPduE*), GS linker, and Golden Gate overhangs | tGGTCTCACatgaataccgacgcaattgaatcgatggtccgggacgtattgagccgcGGCAGCTGAGACCa |
| oCEM208 | Golden Gate cloning | Amplify oCEM207 and oCEM266 - 268 to add buffer regions outside of BsaI cut sites (Fwd) | gggtGGTCTCACatga |
| oCEM209 | Golden Gate cloning | Amplify oCEM207 and oCEM266 - 271 to add buffer regions outside of BsaI cut sites (Rev) | gggtGGTCTCAGCTGC |
| oCEM266 | Golden Gate cloning | Oligo containing *ssPduE*$^{\Delta ESMV::ETLI}$, GS linker, and Golden Gate overhangs | tGGTCTCACatgaataccgacgcaattGAAACTCTTATTcgggacgtattgagccgcGGCAGCTGAGACCa |
| oCEM267 | Golden Gate cloning | Oligo containing *ssPduE*$^{\Delta ESMV::QRIV}$, GS linker, and Golden Gate overhangs | tGGTCTCACatgaataccgacgcaattCAGCGTATTGTAcg |

| | | | |
|---|---|---|---|
| | | | ggacgtattgagccgcGGCAG CTGAGACCa |
| oCEM268 | Golden Gate cloning | Oligo containing *ssPduE*$^{\Delta ESMV::RQII}$, GS linker, and Golden Gate overhangs | tGGTCTCACatgaataccgac gcaattCGTCAGATTATCcg ggacgtattgagccgcGGCAG CTGAGACCa |
| oCEM265 | Golden Gate cloning | Amplify oCEM269 - 271 to add buffer regions outside of BsaI cut sites (Fwd) | gggtGGTCTCACatgg |
| oCEM269 | Golden Gate cloning | Oligo containing *ssPduL*$^{\Delta QSTV::ETLI}$, GS linker, and Golden Gate overhangs | tGGTCTCACatggataaagag cttctgGAAACTCTTATTcgt aaagttctcgacgagGGCAGC TGAGACCa |
| oCEM270 | Golden Gate cloning | Oligo containing *ssPduL*$^{\Delta QSTV::QRIV}$, GS linker, and Golden Gate overhangs | tGGTCTCACatggataaagag cttctgCAGCGTATTGTAcgt aaagttctcgacgagGGCAGC TGAGACCa |
| oCEM271 | Golden Gate cloning | Oligo containing *ssPduL*$^{\Delta QSTV::RQII}$, GS linker, and Golden Gate overhangs | tGGTCTCACatggataaagag cttctgCGTCAGATTATCcgt aaagttctcgacgagGGCAGC TGAGACCa |
| oCEM240 | Golden Gate cloning | Amplify *pduM* to add Golden Gate overhang (Fwd) | AttGGTCTCACATGAACG GCGAAACCCTG |
| oCEM241 | Golden Gate cloning | Amplify *pduM* and *pduM*$^{24-*}$ to add Golden Gate overhang and GS linker (Rev) | ATTGGTCTCAGCTGCCct cctgcttaattaattgaatattccgcg |
| oCEM242 | Golden Gate cloning | Amplify *pduM*$^{24-*}$ to add Golden Gate overhang (Fwd) | AttGGTCTCACATGACGG CGACGCTGAG |

**Supplementary Table S4.** Number of cells counted per replicate for the puncta counts shown in Figures 3b, 4b, and 5b.

| | | Cells Counted per Replicate | | | |
|---|---|---|---|---|---|
| **Genotype** | **Reporter** | **1** | **2** | **3** | **Total** |
| WT | ssPduD-GFP | 78 | 69 | 84 | 231 |
| WT | ssPduP-GFP | 113 | 79 | 66 | 258 |
| WT | ssPduL-GFP | 72 | 142 | 58 | 272 |
| WT | PduG-GFP | 93 | 166 | 106 | 365 |
| WT | PduO-GFP | 160 | 169 | 116 | 445 |
| WT | ssPduM-GFP | 211 | 68 | 90 | 369 |
| WT | PduA-GFP | 72 | 66 | 59 | 197 |
| Δ*ssPduD* | ssPduD-GFP | 69 | 92 | 112 | 273 |
| Δ*ssPduD* | ssPduP-GFP | 86 | 90 | 103 | 279 |
| Δ*ssPduD* | ssPduL-GFP | 82 | 70 | 139 | 291 |
| Δ*ssPduD* | PduG-GFP | 85 | 71 | 47 | 203 |
| Δ*ssPduD* | PduO-GFP | 115 | 118 | 124 | 357 |
| Δ*ssPduD* | PduA-GFP | 96 | 95 | 73 | 264 |

| | | | | | |
|---|---|---|---|---|---|
| Δ*ssPduP* | ssPduD-GFP | 45 | 99 | 88 | 232 |
| Δ*ssPduP* | ssPduP-GFP | 88 | 88 | 74 | 250 |
| Δ*ssPduP* | ssPduL-GFP | 82 | 130 | 147 | 359 |
| Δ*ssPduP* | PduG-GFP | 110 | 118 | 92 | 320 |
| Δ*ssPduP* | PduO-GFP | 142 | 65 | 91 | 298 |
| Δ*ssPduP* | PduA-GFP | 139 | 44 | 92 | 275 |
| Δ*ssPduL* | ssPduD-GFP | 77 | 84 | 146 | 307 |
| Δ*ssPduL* | ssPduP-GFP | 72 | 72 | 126 | 270 |
| Δ*ssPduL* | ssPduL-GFP | 94 | 82 | 76 | 252 |
| Δ*ssPduL* | PduG-GFP | 57 | 59 | 84 | 200 |
| Δ*ssPduL* | PduO-GFP | 112 | 73 | 88 | 273 |
| Δ*ssPduL* | PduA-GFP | 143 | 155 | 122 | 420 |
| Δ*ssPduDP* | ssPduD-GFP | 77 | 105 | 66 | 248 |
| Δ*ssPduDP* | ssPduP-GFP | 71 | 117 | 117 | 305 |
| Δ*ssPduDP* | ssPduL-GFP | 100 | 120 | 157 | 377 |
| Δ*ssPduDP* | PduG-GFP | 67 | 56 | 146 | 269 |
| Δ*ssPduDP* | PduO-GFP | 70 | 131 | 124 | 325 |
| Δ*ssPduDP* | PduA-GFP | 163 | 66 | 59 | 288 |
| Δ*ssPduDL* | ssPduD-GFP | 132 | 69 | 64 | 265 |
| Δ*ssPduDL* | ssPduP-GFP | 83 | 74 | 148 | 305 |
| Δ*ssPduDL* | ssPduL-GFP | 88 | 101 | 86 | 275 |
| Δ*ssPduDL* | PduG-GFP | 91 | 173 | 83 | 347 |
| Δ*ssPduDL* | PduO-GFP | 81 | 110 | 115 | 306 |
| Δ*ssPduDL* | PduA-GFP | 188 | 94 | 72 | 354 |
| Δ*ssPduPL* | ssPduD-GFP | 74 | 97 | 114 | 285 |
| Δ*ssPduPL* | ssPduP-GFP | 92 | 93 | 138 | 323 |
| Δ*ssPduPL* | ssPduL-GFP | 105 | 62 | 76 | 243 |
| Δ*ssPduPL* | PduG-GFP | 75 | 136 | 99 | 310 |
| Δ*ssPduPL* | PduO-GFP | 70 | 65 | 96 | 231 |
| Δ*ssPduPL* | PduA-GFP | 130 | 127 | 113 | 370 |
| Δ*ssPduDPL* | ssPduD-GFP | 115 | 89 | 113 | 317 |
| Δ*ssPduDPL* | ssPduP-GFP | 79 | 73 | 119 | 271 |
| Δ*ssPduDPL* | ssPduL-GFP | 100 | 93 | 57 | 250 |
| Δ*ssPduDPL* | PduG-GFP | 95 | 54 | 98 | 247 |
| Δ*ssPduDPL* | PduO-GFP | 99 | 81 | 134 | 314 |
| Δ*ssPduDPL* | PduA-GFP | 104 | 108 | 61 | 273 |
| Δ*pduB* | ssPduD-GFP | 135 | 214 | 275 | 624 |
| Δ*pduB* | ssPduM-GFP | 43 | 104 | 111 | 258 |
| Δ*pduB* | PduG-GFP | 232 | 311 | 233 | 776 |
| Δ*pduB* | PduA-GFP | 40 | 33 | 179 | 252 |
| Δ*ssPduB* | ssPduD-GFP | 110 | 60 | 172 | 342 |
| Δ*ssPduB* | ssPduM-GFP | 268 | 108 | 108 | 484 |
| Δ*ssPduB* | PduG-GFP | 166 | 53 | 57 | 276 |
| Δ*ssPduB* | PduA-GFP | 82 | 124 | 137 | 343 |
| Δ*pduM* | ssPduD-GFP | 231 | 205 | 216 | 652 |

| | | | | | |
|---|---|---|---|---|---|
| Δ*pduM* | ssPduM-GFP | 129 | 168 | 221 | 518 |
| Δ*pduM* | PduG-GFP | 159 | 167 | 157 | 483 |
| Δ*pduM* | PduA-GFP | 149 | 111 | 60 | 320 |
| Δ*ssPduM* | ssPduD-GFP | 96 | 42 | 108 | 246 |
| Δ*ssPduM* | ssPduM-GFP | 80 | 92 | 178 | 350 |
| Δ*ssPduM* | PduG-GFP | 238 | 119 | 136 | 493 |
| Δ*ssPduM* | PduA-GFP | 33 | 50 | 58 | 141 |
| Δ*ssPduMB* | ssPduD-GFP | 55 | 76 | 100 | 231 |
| Δ*ssPduMB* | ssPduM-GFP | 152 | 52 | 109 | 313 |
| Δ*ssPduMB* | PduG-GFP | 81 | 159 | 55 | 295 |
| Δ*ssPduMB* | PduA-GFP | 68 | 46 | 106 | 220 |
| Δ*ssPduDPLB* | ssPduD-GFP | 79 | 72 | 127 | 278 |
| Δ*ssPduDPLB* | ssPduM-GFP | 37 | 113 | 47 | 197 |
| Δ*ssPduDPLB* | PduG-GFP | 80 | 84 | 93 | 257 |
| Δ*ssPduDPLB* | PduA-GFP | 59 | 46 | 54 | 159 |
| Δ*ssPduDPLM* | ssPduD-GFP | 103 | 109 | 82 | 294 |
| Δ*ssPduDPLM* | ssPduM-GFP | 90 | 45 | 91 | 226 |
| Δ*ssPduDPLM* | PduG-GFP | 230 | 93 | 161 | 484 |
| Δ*ssPduDPLM* | PduA-GFP | 108 | 97 | 88 | 293 |
| Δ*ssPduDPLMB* | ssPduD-GFP | 58 | 131 | 132 | 321 |
| Δ*ssPduDPLMB* | ssPduM-GFP | 73 | 117 | 88 | 278 |
| Δ*ssPduDPLMB* | PduG-GFP | 105 | 140 | 163 | 408 |
| Δ*ssPduDPLMB* | PduA-GFP | 86 | 178 | 208 | 472 |

**Supplementary Table S5.** Designs and results of statistical tests conducted in this study.

| Test description | One-factor ANOVA between puncta counts of core reporters expressed in a strain, Bonferroni post-hoc test |
|---|---|
| Reporters included in test | ssPduD-GFP, ssPduP-GFP, ssPduL-GFP, PduG-GFP, PduO-GFP |
| *F* statistic and *p* value, Δ*ssPduD* | $F = 129.08$, $p = 1.462\times10^{-8}$ |
| *F* statistic and *p* value, Δ*ssPduP* | $F = 9.41$, $p = 0.0020$ |
| *F* statistic and *p* value, Δ*ssPduL* | $F = 7.67$, $p = 0.0043$ |
| *F* statistic and *p* value, Δ*ssPduDP* | $F = 183.13$, $p = 2.628\times10^{-9}$ |
| *F* statistic and *p* value, Δ*ssPduDL* | $F = 104.58$, $p = 4.080\times10^{-8}$ |
| *F* statistic and *p* value, Δ*ssPduPL* | $F = 4.18$, $p = 0.0303$ |

| | |
|---|---|
| *F* statistic and *p* value, Δ*ssPduDPL* | $F$ = 27.41, $p$ = 2.278×10$^{-5}$ |
| Total degrees of freedom | 14 (Between groups df = 4, within groups df = 10) |
| Test description | One-factor ANOVA between puncta counts of different reporters expressed in a strain, Dunnett post-hoc test |
| Control group | PduA-GFP |
| Experimental groups | ssPduD-GFP, ssPduM-GFP, PduG-GFP |
| *F* statistic and *p* value, Δ*pduM* | $F$ = 21.94, $p$ = 3.25×10$^{-4}$ |
| *F* statistic and *p* value, Δ*ssPduM* | $F$ = 25.15, $p$ = 2.00×10$^{-4}$ |
| *F* statistic and *p* value, Δ*ssPduDPLB* | $F$ = 37.37, $p$ = 4.71×10$^{-5}$ |
| *F* statistic and *p* value, Δ*ssPduDPLMB* | $F$ = 36.12, $p$ = 5.35×10$^{-5}$ |
| Total degrees of freedom | 11 (Between groups df = 3, within groups df = 8) |
| Control group | ssPduD-GFP |
| Experimental groups | ssPduM-GFP, PduG-GFP |
| *F* statistic and *p* value, Δ*ssPduDPLB* | $F$ = 0.38, $p$ = 0.70 |
| *F* statistic and *p* value, Δ*ssPduDPLM* | $F$ = 0.76, $p$ = 0.51 |
| *F* statistic and *p* value, Δ*ssPduDPLMB* | $F$ = 1.62, $p$ = 0.27 |
| Total degrees of freedom | 8 (Between groups df = 2, within groups df = 6) |
| Test description | One-factor ANOVA between puncta counts of PduA-GFP expressed in different strains, Bonferroni post-hoc test |
| Strains included in test | WT, Δ*pduB*, Δ*ssPduB*, Δ*pduM*, Δ*ssPduM*, Δ*ssPduMB*, Δ*ssPduDPLB*, Δ*ssPduDPLM*, Δ*ssPduDPLMB* |
| *F* statistic and *p* value | $F$ = 26.32, $p$ = 2.06×10$^{-8}$ |
| Total degrees of freedom | 26 (Between groups df = 8, within groups df = 18) |
| Test description | Two-factor ANOVA between puncta counts of reporters expressed in two strains |
| Strains included in test | Δ*ssPduM*, Δ*pduM* |
| Reporters included in test | ssPduD-GFP, ssPduM-GFP, PduG-GFP, PduA-GFP |
| *F* statistic and *p* value, strains | $F$ = 38.96, $p$ = 1.18×10$^{-5}$ |
| *F* statistic and *p* value, reporters | $F$ = 38.24, $p$ = 1.59×10$^{-7}$ |

| | |
|---|---|
| *F* statistic and *p* value, interaction | $F = 7.17$, $p = 0.0029$ |
| Total degrees of freedom | 23 (Between strains df = 1, between reporters df = 3, interaction df = 3, within groups df = 16) |
| Simple main effects test description | Two-tailed Student's *t*-test between strains for each reporter |
| *t* statistic and *p* value, ssPduD-GFP | $t = 12.31$, $p = 2.50\times10^{-4}$ |
| *t* statistic and *p* value, ssPduM-GFP | $t = 2.21$, $p = 0.091$ |
| *t* statistic and *p* value, PduG-GFP | $t = 0.76$, $p = 0.49$ |
| *t* statistic and *p* value, PduA-GFP | $t = 3.06$, $p = 0.0376$ |
| Degrees of freedom, simple main effects *t*-tests | 4 |

**Supplementary Table S6.** Bonferroni post hoc test from ANOVA used in Figure 3b. This table shows pairwise *p*-values between normalized puncta counts of enzymatic signal sequences and other core reporters in enzymatic signal sequence knockout strains.

| Reporter 1 | Reporter 2 | *p*, Δ*ssD* | *p*, Δ*ssP* | *p*, Δ*ssL* | *p*, Δ*ssDP* | *p*, Δ*ssDL* | *p*, Δ*ssPL* | *p*, Δ*ssDPL* |
|---|---|---|---|---|---|---|---|---|
| ssPduD-GFP | ssPduP-GFP | $8.65\times10^{-7}$ | 1 | 1 | $2.01\times10^{-7}$ | $1.33\times10^{-7}$ | 1 | $6.84\times10^{-5}$ |
| ssPduD-GFP | ssPduL-GFP | $8.14\times10^{-7}$ | 1 | 0.3579 | $8.76\times10^{-9}$ | $1.56\times10^{-6}$ | 0.130 | $6.25\times10^{-3}$ |
| ssPduD-GFP | PduG-GFP | $2.32\times10^{-8}$ | 1 | 1 | $9.16\times10^{-9}$ | $6.03\times10^{-8}$ | 1 | $2.85\times10^{-5}$ |
| ssPduD-GFP | PduO-GFP | $2.77\times10^{-8}$ | $4.41\times10^{-3}$ | 0.1545 | $6.75\times10^{-9}$ | $3.57\times10^{-7}$ | 1 | $4.54\times10^{-3}$ |
| ssPduP-GFP | ssPduL-GFP | 1 | 1 | 0.0320 | $1.28\times10^{-3}$ | 0.0379 | 1 | 0.0477 |
| ssPduP-GFP | PduG-GFP | $1.03\times10^{-3}$ | 1 | 1 | $1.46\times10^{-3}$ | 1 | 1 | 1 |
| ssPduP-GFP | PduO-GFP | $1.64\times10^{-3}$ | $6.93\times10^{-3}$ | 0.0146 | $5.96\times10^{-4}$ | 1 | 0.429 | 0.0687 |
| ssPduL-GFP | PduG-GFP | $1.15\times10^{-3}$ | 1 | 0.0795 | 1 | $4.24\times10^{-3}$ | 0.6156 | 0.0118 |
| ssPduL-GFP | PduO-GFP | $1.85\times10^{-3}$ | 0.0322 | 1 | 1 | 0.582 | 0.0393 | 1 |
| PduG-GFP | PduO-GFP | 1 | $5.07\times10^{-3}$ | 0.0352 | 1 | 0.1284 | 1 | 0.0166 |

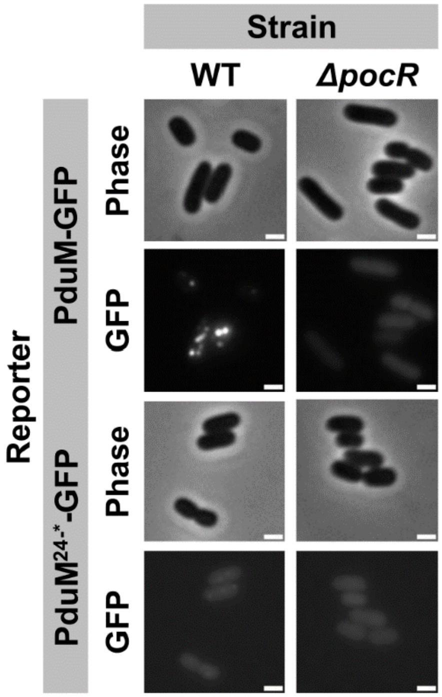


**Supplementary Figure S1.** Optical and fluorescence micrographs of full length PduM and PduM$^{24-*}$ (PduM without its signal sequence) fused to GFPmut2. These constructs were overexpressed both in wild-type (WT) MCP-forming *S. enterica* and in Δ*pocR*, an assembly-deficient *S. enterica* strain in which all MCP formation is abolished. All scale bars are 1 µm. Similar results were observed across at least three biological replicates of each strain.

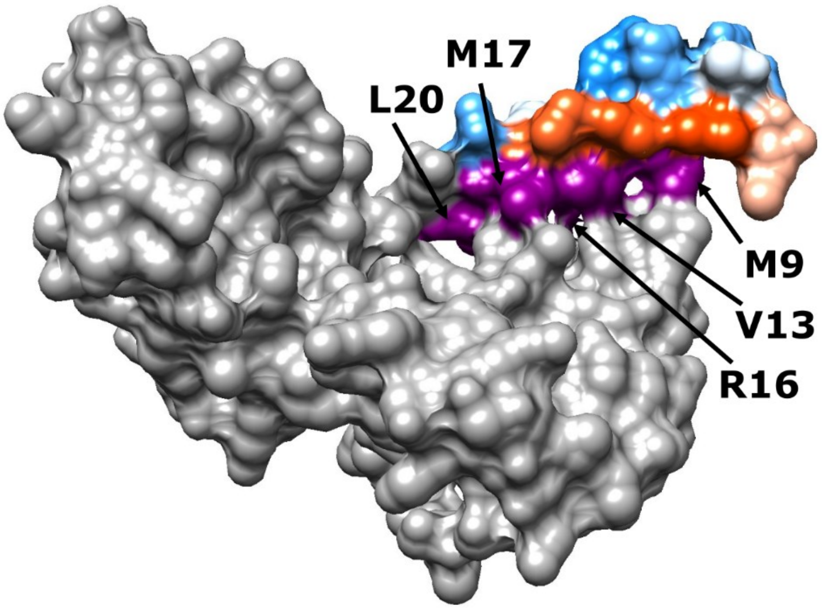


**Supplementary Figure S2.** Predicted hydrophobicity surface of PduE from *S. enterica* LT2. The body of PduE is colored gray, and the signal sequence-like N-terminal motif (ssPduE) is colored in blue, red, and purple. Hydrophilic areas are shown in blue, hydrophobic areas are shown in red, and residues that are predicted to connect ssPduE to the body of PduE are shown in purple and labeled with arrows. This structure was downloaded from the AlphaFold Protein Structure Database and visualized using UCSF Chimera.[1-3]

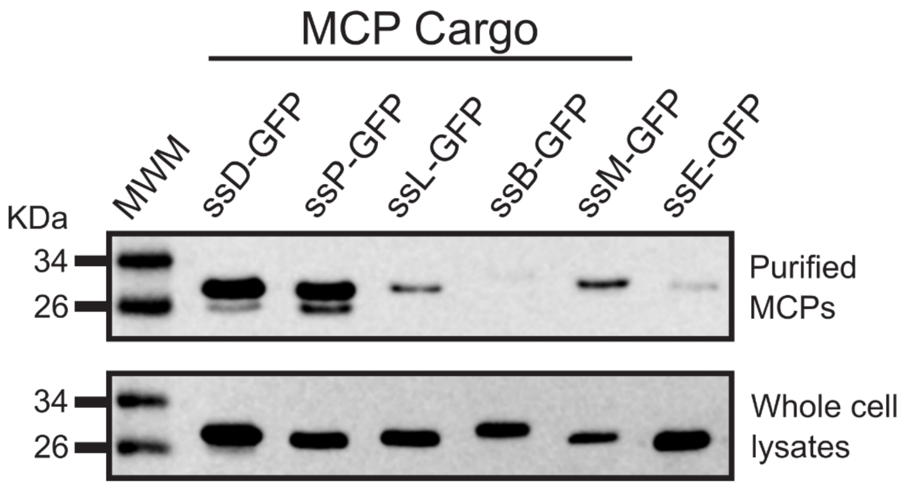


**Supplementary Figure S3.** Anti-GFP western blots of purified MCPs and whole cell lysates from wild-type *Salmonella enterica* overexpressing each signal sequence fused to GFPmut2. Similar results were observed across two independent biological replicates.

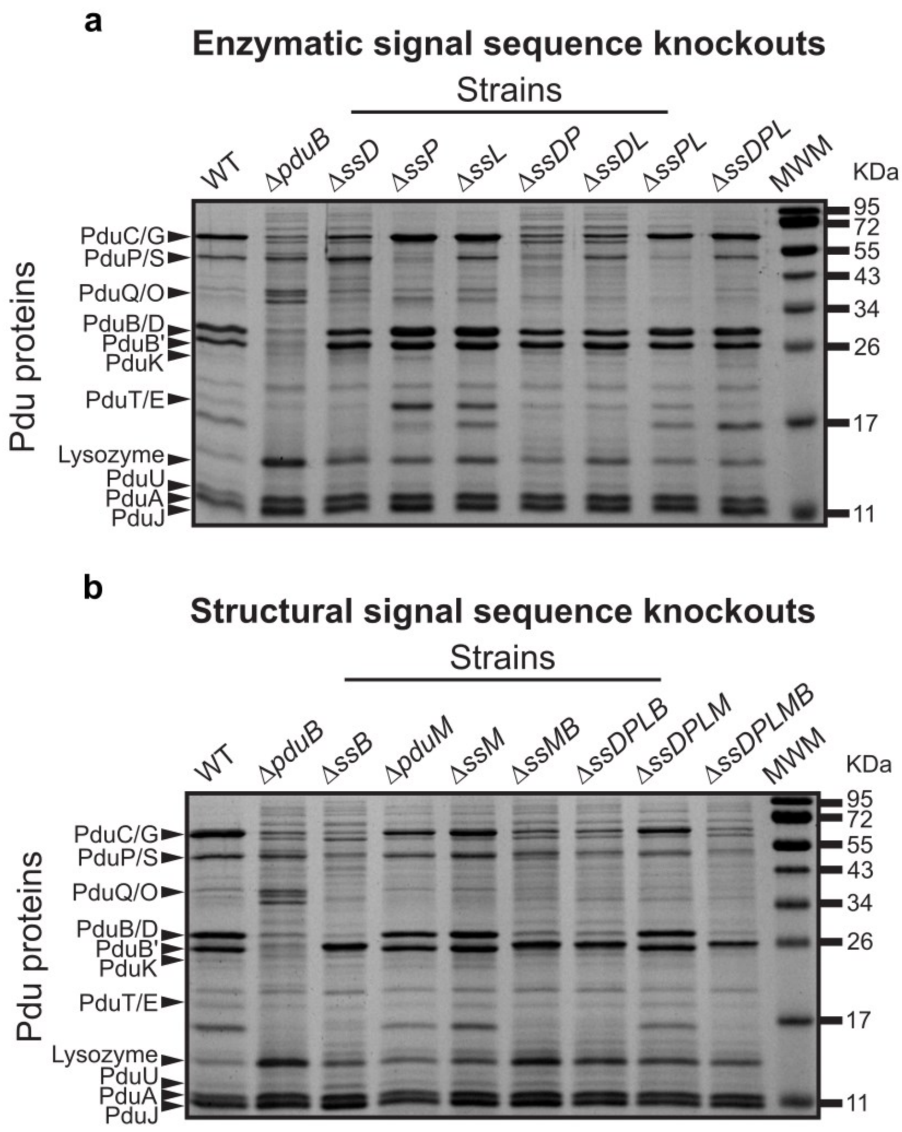


**Supplementary Figure S4.** Coomassie stained SDS-PAGE of MCPs purified from (a) enzymatic signal sequence knockout strains and (b) structural and structural + enzymatic signal sequence knockout strains. Bands corresponding to various Pdu proteins and lysozyme are labeled, and molecular weight markers (MWM) are included on the right side of the gels. Similar results were observed across three technical replicates.

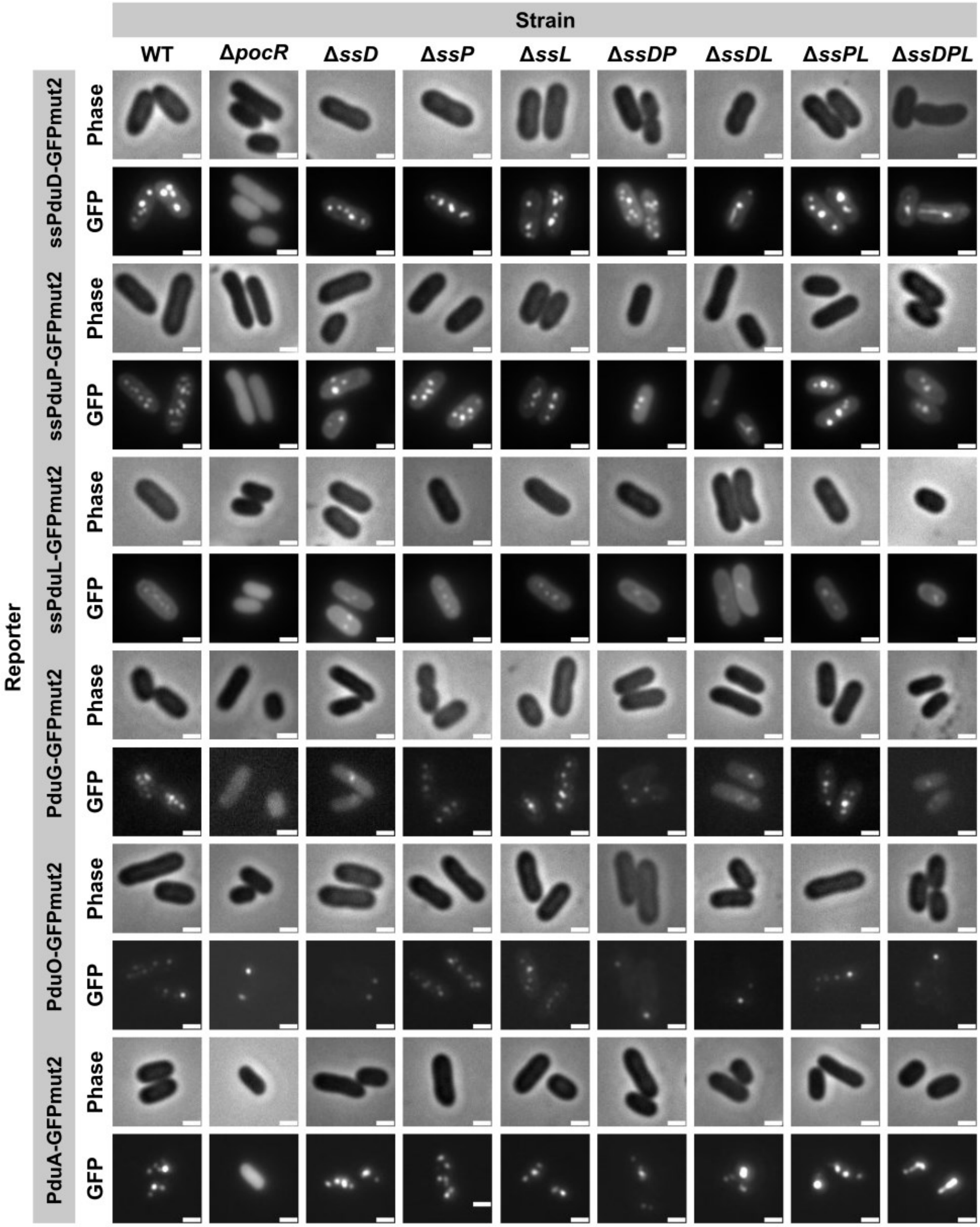


**Supplementary Figure S5.** Optical and fluorescence micrographs of Pdu MCP core and shell proteins and signal sequences fused to GFPmut2. These constructs were expressed in wild type (WT) *S. enterica*, in Δ*pocR*, an assembly-deficient strain in which all MCP formation is abolished, and in the enzymatic signal sequence knockout strains. All scale bars are 1 μm. Similar results were observed across at least three biological replicates of each strain.

**a**

| | WT | Δ*ssD* | Δ*ssP* | Δ*ssL* | Δ*ssDP* | Δ*ssDL* | Δ*ssPL* | Δ*ssDPL* |
|---|---|---|---|---|---|---|---|---|
| ssD-GFP | 3.7 ± 0.3 | 4.0 ± 0.2 | 3.3 ± 0.2 | 2.74 ± 0.09 | 3.97 ± 0.08 | 2.47 ± 0.09 | 2.7 ± 0.1 | 1.75 ± 0.07 |
| ssP-GFP | 4.9 ± 0.2 | 2.8 ± 0.4 | 4.6 ± 0.01 | 3.3 ± 0.2 | 2.6 ± 0.3 | 1.47 ± 0.09 | 3.3 ± 0.1 | 1.3 ± 0.2 |
| ssL-GFP | 3.1 ± 0.2 | 1.72 ± 0.04 | 3.0 ± 0.3 | 2.7 ± 0.3 | 1.0 ± 0.1 | 1.18 ± 0.06 | 1.9 ± 0.1 | 1.10 ± 0.03 |
| PduG-GFP | 4.0 ± 0.5 | 1.3 ± 0.1 | 3.63 ± 0.06 | 2.8 ± 0.2 | 1.3 ± 0.3 | 1.1 ± 0.2 | 2.8 ± 0.1 | 1.00 ± 0.03 |
| PduO-GFP | 3.3 ± 0.1 | 1.09 ± 0.09 | 3.9 ± 0.3 | 2.9 ± 0.1 | 1.00 ± 0.03 | 1.10 ± 0.05 | 2.5 ± 0.3 | 1.2 ± 0.1 |
| PduA-GFP | 3.8 ± 0.4 | 3.1 ± 0.4 | 4.0 ± 0.6 | 2.9 ± 0.1 | 2.7 ± 0.1 | 1.8 ± 0.1 | 2.5 ± 0.1 | 2.1 ± 0.1 |

**b**

| | WT | Δ*pduB* | Δ*ssB* | Δ*pduM* | Δ*ssM* | Δ*ssMB* | Δ*ssDPLB* | Δ*ssDPLM* | Δ*ssDPLMB* |
|---|---|---|---|---|---|---|---|---|---|
| ssD-GFP | 3.7 ± 0.3 | 0.92 ± 0.04 | 1.1 ± 0.1 | 2.4 ± 0.1 | 1.52 ± 0.03 | 1.00 ± 0.02 | 1.05 ± 0.08 | 1.4 ± 0.2 | 0.82 ± 0.08 |
| PduG-GFP | 4.0 ± 0.5 | 1.01 ± 0.02 | 0.99 ± 0.04 | 1.81 ± 0.06 | 1.7 ± 0.2 | 0.97 ± 0.04 | 1.04 ± 0.04 | 1.6 ± 0.1 | 0.90 ± 0.10 |
| ssM-GFP | 4.7 ± 0.4 | 0.94 ± 0.01 | 0.96 ± 0.02 | 2.04 ± 0.07 | 1.9 ± 0.1 | 0.98 ± 0.03 | 1.01 ± 0.05 | 1.5 ± 0.3 | 0.94 ± 0.05 |
| PduA-GFP | 3.8 ± 0.4 | 3.6 ± 0.3 | 3.3 ± 0.1 | 3.3 ± 0.5 | 2.5 ± 0.2 | 3.1 ± 0.2 | 2.0 ± 0.3 | 1.5 ± 0.1 | 1.8 ± 0.2 |

**Supplementary Figure S6.** Means and standard deviations of GFP puncta per cell for (a) strains and reporters shown in the main figure 3b heatmap and (b) strains and reporters shown in the main figures 4b and 5b heatmaps. The values shown in this figure are the means and standard deviations over three biological replicates of at least 30 cells each and are not normalized to wild type puncta counts.

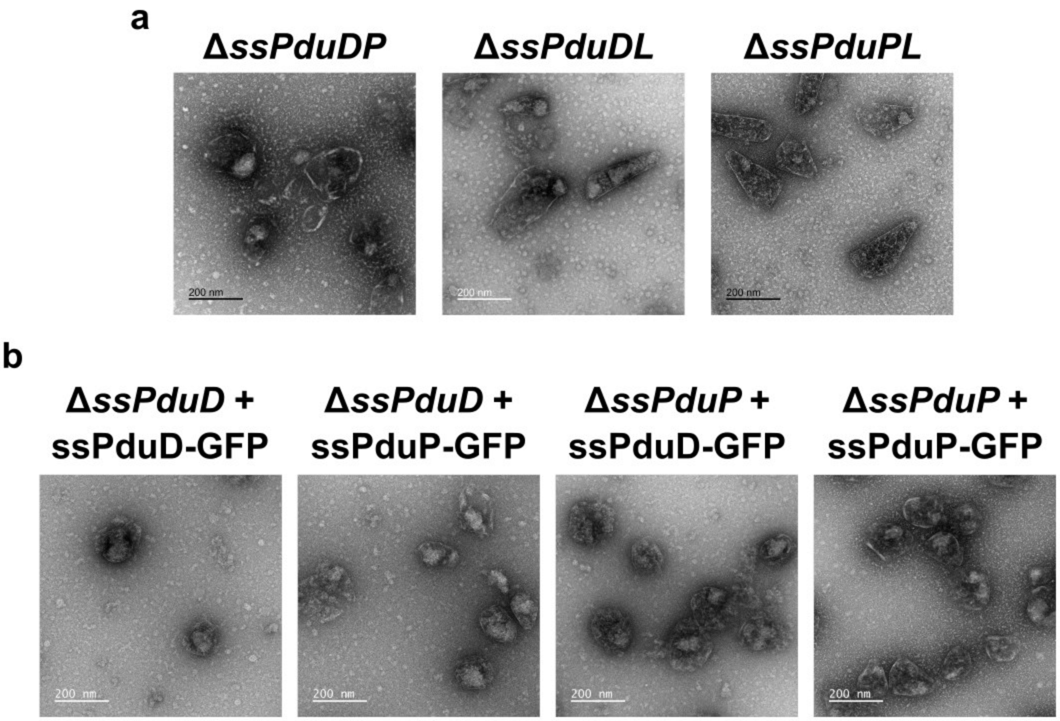


**Supplementary Figure S7.** Transmission electron micrographs of purified MCPs from (a) strains with two enzymatic signal sequences knocked out and (b) Δ*ssPduD* and Δ*ssPduP* complemented with overexpressed ssPduD-GFP and ssPduP-GFP. These images are representative of multiple images taken of the same sample, but due to time constraints, cost constraints, and a large number of samples, transmission electron micrographs were only taken of one biological replicate.

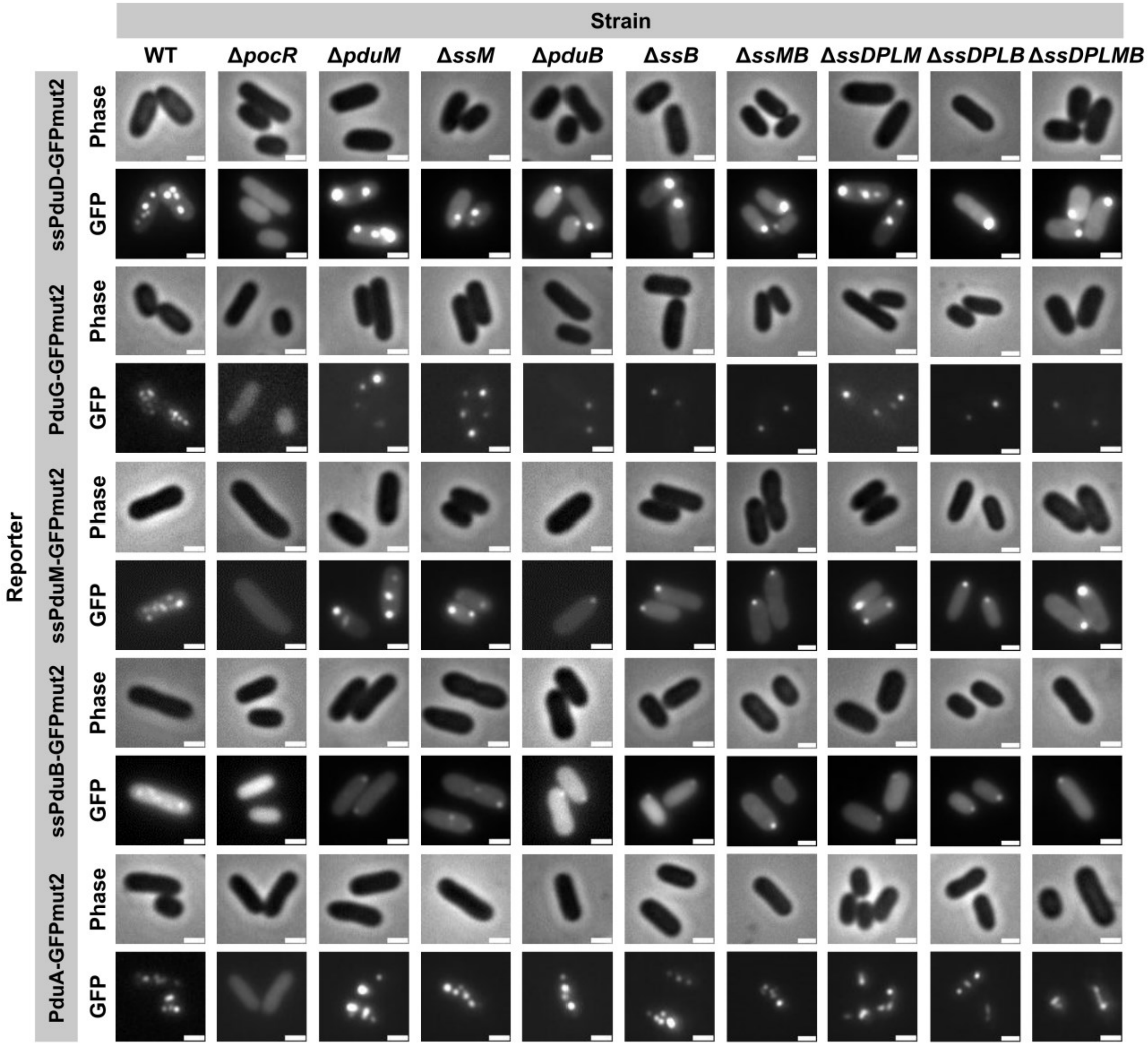


**Supplementary Figure S8.** Optical and fluorescence micrographs of Pdu MCP core and shell proteins and signal sequences fused to GFPmut2. These constructs were expressed in wild type (WT) *S. enterica*, in Δ*pocR*, an assembly-deficient strain in which all MCP formation is abolished, and in the structural and enzymatic + structural signal sequence knockout strains. All scale bars are 1 μm. Similar results were observed across at least three biological replicates of each strain.

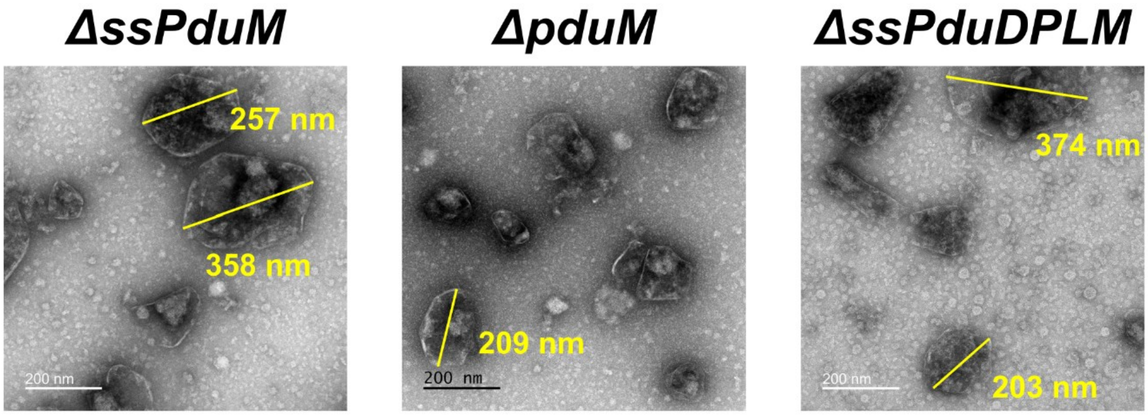


**Supplementary Figure S9.** Transmission electron micrographs of purified Δ*ssPduM*, Δ*pduM*, and Δ*ssPduDPLM* MCPs noting the sizes of MCPs over 200 nm in diameter. These images are representative of multiple images taken of the same sample, but due to time constraints, cost constraints, and a large number of samples, transmission electron micrographs were taken of only one biological replicate.
